# Discovery of a *Pseudomonas aeruginosa* Type VI secretion system toxin targeting bacterial protein synthesis using a global genomics approach

**DOI:** 10.1101/733030

**Authors:** Laura M. Nolan, Amy K. Cain, Eleni Manoli, Maria A. Sainz-Polo, Gordon Dougan, Despoina A.I. Mavridou, David Albesa-Jové, Julian Parkhill, Alain Filloux

**Affiliations:** MRC Centre for Molecular Bacteriology and Infection (CMBI), Department of Life Sciences, Imperial College London, London SW7 2AZ, United Kingdom; Wellcome Trust Sanger Institute, Wellcome Trust Genome Campus, Hinxton, Cambridge, United Kingdom; Structural Biology Unit, CIC bioGUNE, Bizkaia Technology Park, 48160 Derio, Spain; IKERBASQUE, Basque Foundation for Science, Bilbao, Spain

## Abstract

The Type VI secretion system (T6SS) is a bacterial weapon which delivers toxic effectors to kill competitors or subvert some of their key functions. Here we use <u>tra</u>nsposon <u>d</u>irected insertion-site <u>s</u>equencing (TraDIS) to identify T6SS toxins associated with the H1-T6SS, one of the three T6SS machines found in *Pseudomonas aeruginosa*. This approach identified several putative toxin-immunity pairs, including Tse8-Tsi8. Full characterization of this protein pair demonstrated that Tse8 is delivered by the VgrG1a spike complex into prey cells where it targets the transamidosome, a multiprotein complex involved in protein synthesis in bacteria lacking either one or both of the asparagine or glutamine tRNA synthases. Our data suggests that Tse8 combines as a non-cognate component of the transamidosome complex, reducing fitness by limiting the ability of the cell to synthesize proteins. This is the first demonstration of a T6SS toxin affecting protein synthesis, expanding the range of cellular components targeted by this bacterial weapon. The success of the current study validates the use of our TraDIS approach as a tool to drastically expand the repertoire of T6SS toxins in any T6SS-encoding bacterium.

---

Bacteria rarely exist in a single-species planktonic state and instead form complex polymicrobial structures, called biofilms^1, 2^. Within this context bacteria often compete with other microorganisms to secure space and nutrients. The Type VI secretion system (T6SS) is a Gram-negative bacterial weapon which delivers toxins into neighbouring competitors to either kill or subvert their key functions in order to attain dominance within a given niche^3–5^. The T6SS is composed of 13 core components, several of which are structurally related to proteins from the T4 bacteriophage tail^6^. The Hcp tube-like structure is capped by a VgrG-PAAR tip complex, or spike, and encapsulated within a TssBC, or VipAB, contractile sheath. Upon extension of the sheath within the cytoplasm and subsequent contraction, the spike is thought to facilitate the puncturing of the cell membranes of both the producing and target cells, allowing delivery of the attached toxins^7, 8^. T6SS toxins have been shown to be secreted in association with the VgrG tip complex, the Hcp tube, or as extension domains of the VgrG, PAAR or Hcp proteins^9–12^. Importantly, neighbouring bacterial sister cells are protected from the effects of the toxins by production of cognate immunity proteins, which are usually encoded adjacent to the toxin gene in the genome^13^. The major identified targets of T6SS toxins to date are components of the cell wall, as well as the cell membrane and nucleic acids^14^. These T6SS toxins have mainly been identified by searching in the genomic proximity of known T6SS components, or by detection of toxins in the secretome^9, 12, 15^.

*Pseudomonas aeruginosa* is a highly antibiotic-resistant Gram-negative pathogen and ranked second by the World Health Organization in the list of bacteria that require immediate attention. It is also a highly potent T6SS bacterial killer, equipped with three independent systems (H1-to H3-T6SS)^16^. In the current study we used a global genomics-based approach called TraDIS (<u>Tra</u>nsposon <u>d</u>irected insertion-site <u>s</u>equencing) to identify novel toxins associated with the *P. aeruginosa* H1-T6SS. A previous study has used Tn-Seq, a similar global transposon mutagenesis approach, and confirmed the presence of three T6SS toxin-immunity genes which are located in the vicinity of *vgrG* genes in *V. cholerae*^41^. Our TraDIS approach identified several remote and novel putative T6SS toxin-immunity pairs. We found that one of the identified toxins, Tse8 (<u>T</u>ype <u>s</u>ix <u>e</u>xported 8), targets the bacterial transamidosome complex, which is required for protein synthesis in bacteria that lack the asparagine and/or glutamine tRNA synthases^17^. This is an entirely new target for a T6SS toxin and the first shown to target bacterial protein synthesis.

## TraDIS identifies known and novel H1-T6SS toxin-immunity pairs

To systematically identify *P. aeruginosa* PAK H1-T6SS associated immunity genes we generated duplicate high-density insertion transposon mutant libraries consisting of ∼2 million mutants in a H1-T6SS active (PAKΔ*retS*) and a H1-T6SS inactive (PAKΔ*retS*Δ*H1*) background. We reasoned that transposon insertions in immunity genes would only be tolerated in the H1-T6SS inactive library, while in the H1-T6SS active library, cells lacking an immunity protein would be killed upon injection of the cognate toxin from neighbouring sister cells or due to self-intoxication. Each duplicate library was plated separately at high-contact density on agar plates and passaged in an overnight incubation step to promote T6SS-mediated killing of mutants with transposon insertions in immunity genes (Fig. 1). The genomic DNA of mutants which were not killed in both the H1-T6SS active and inactive libraries were then separately sequenced using a mass-parallel approach as described previously^18, 19^ (Fig. 1). The relative frequencies of transposon insertion in genes in the H1-T6SS active and inactive libraries revealed a large number of genes which had changes in relative numbers of transposon insertions. Forty-five genes which had a significantly greater number of normalized transposon insertions in the H1-T6SS inactive library background, compared to the H1-T6SS active library background, were identified (Supplementary Table 1), and may encode putative H1-T6SS immunity proteins. Our approach is validated by our ability to identify five (*tsi1-tsi5*) out of the seven known H1-T6SS immunity genes, whose gene products protect against cognate toxins acting in both the cytoplasm and periplasm (Table 1). Our screen was unable to identify *tsi6* as this gene is deleted in our PAKΔ*retS*Δ*H1* strain, thus there is no possibility to assess the relative frequency of transposon insertions in this gene between the two library backgrounds. In the case of *tsi7* we did not see any difference in the levels of insertions between the two libraries (Supplementary Table 1). This is probably due to the fact that we also saw insertions in the PAAR domain of its cognate toxin Tse7 in both library backgrounds. Insertions in the PAAR domain are expected to destabilise the interaction of Tse7 with VgrG1b which has been shown to be mediated by the Tse7 PAAR domain^20^ abrogating the activity of the toxin in both the active and inactive libraries.

**Figure 1.**
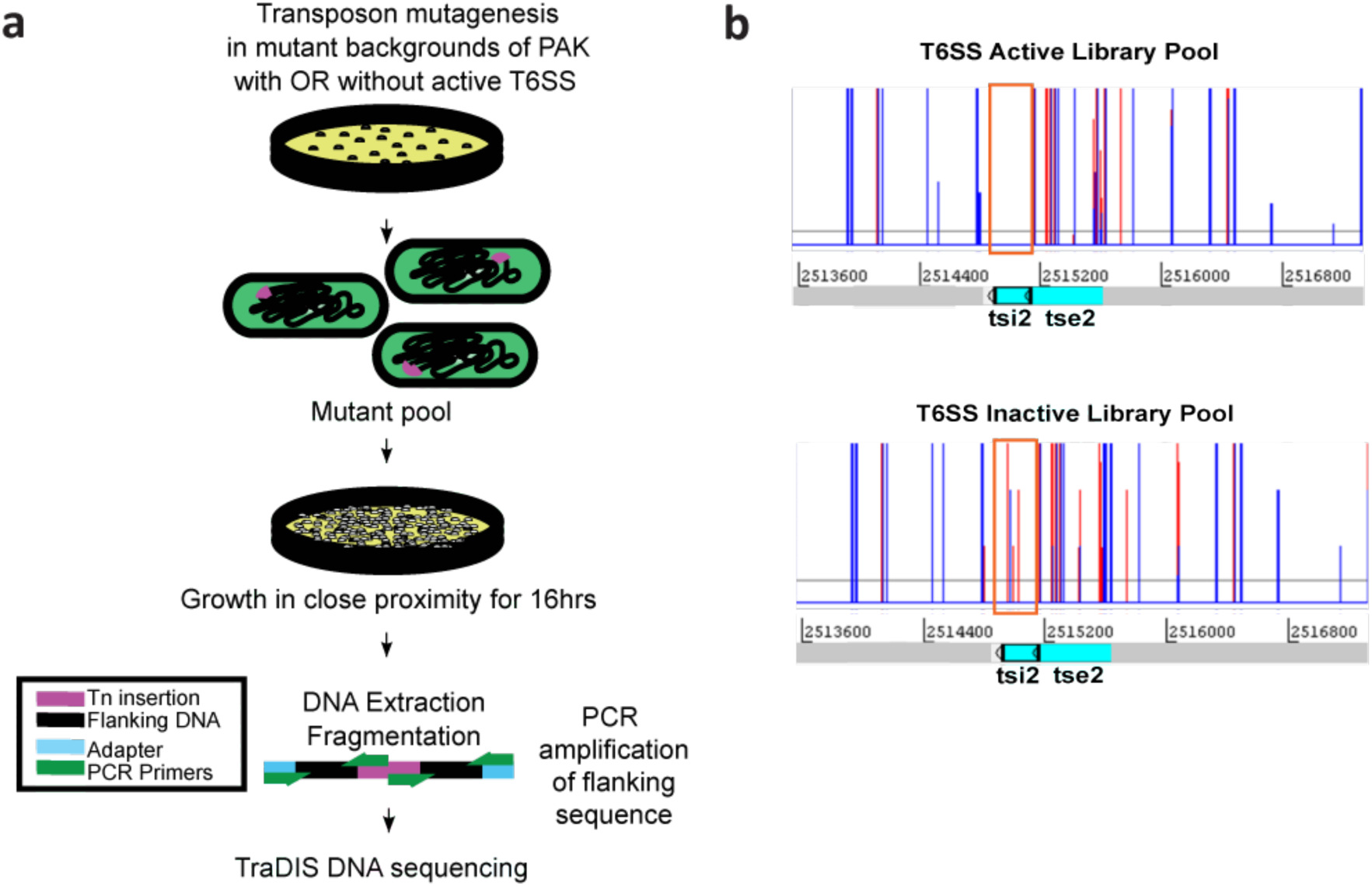
TraDIS library generation and sequencing workflow (a) and predicted outcome of transposon insertions in *tsi* (immunity) genes in each library background (b). **a**, *En masse* transposon (Tn) mutagenesis in T6SS active (PAKΔ*retS*) or T6SS inactive (PAKΔ*retS*ΔH1) backgrounds was performed to generate pooled transposon mutant libraries of ∼2 million mutants each. These libraries were then separately passaged overnight at high contact density and the genomic DNA from recovered mutants was harvested. This genomic DNA was then fragmented and adapters ligated to each end prior to PCR enrichment for transposon-containing DNA fragments. The pooled DNA population was then subjected to TraDIS DNA sequencing. **b,** Artemis (http://www.sanger.ac.uk/science/tools/artemis) plot file showing distribution of transposon insertions (red and blue lines correspond to insertions mapped from either forward or reverse sequence reads) in immunity gene (*tsi2* in this case) in the T6SS active library background (top panel - no insertions permitted) and in the T6SS inactive library background (right - insertions are permitted). The other H1-T6SS immunity genes detected, as well as the putative novel T6SS immunity genes (Table 1) had a similar distribution of transposon insertions in each library background as for *tsi2*. Panel (a) adapted from Barquist *et al.,* (2013)^51^.

**Table 1.**
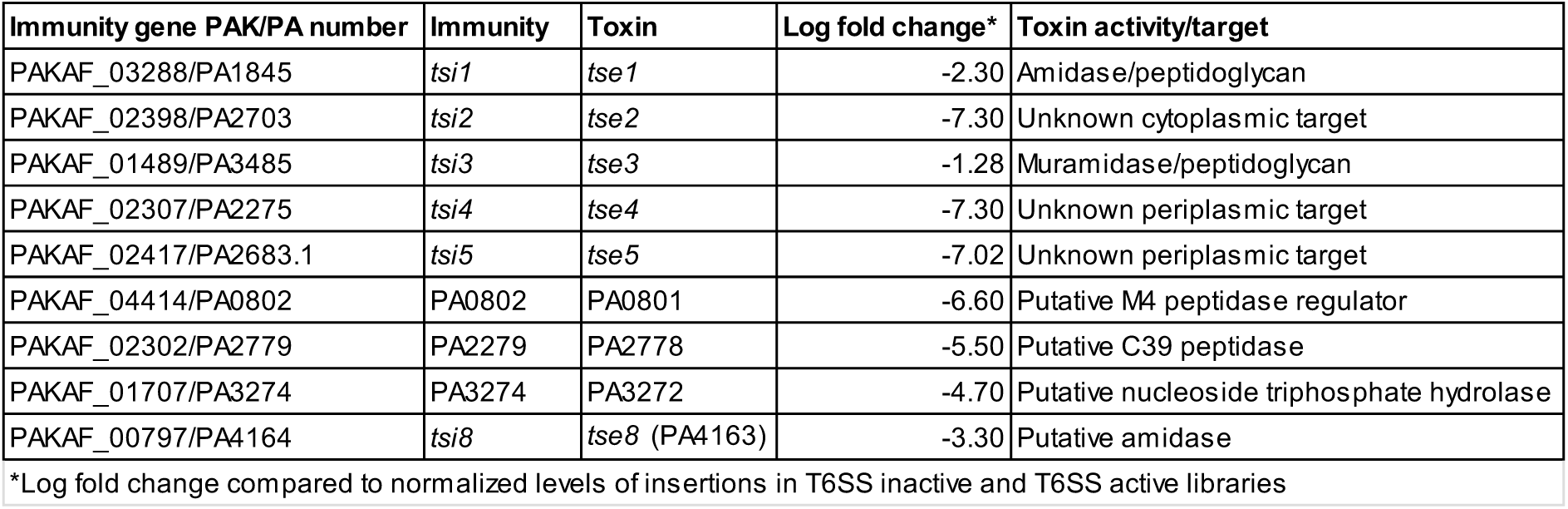
TraDIS allows identification of known and putative novel H1-T6SS immunity genes

In addition to known H1-T6SS associated immunity genes, our TraDIS approach identified multiple uncharacterised small coding sequences, which displayed a decrease in transposon insertions in the H1-T6SS active compared to the inactive background (represented by a negative log fold change), suggesting a role for these genes in protecting against H1-T6SS mediated killing (Supplementary Table 1). Upstream of several of these were genes encoding proteins with putative enzymatic activity which could be T6SS toxins: PAKAF_04413 (PA0801) encodes a putative M4 peptidase regulator; PAKAF_02301 (PA2778) encodes a putative C39 peptidase domain-containing protein; PAKAF_01705 (PA3272) encodes a putative nucleoside triphosphate hydrolase; and PAKAF_00796 (PA4163) encodes a putative amidase (Table 1 and Extended Data Fig. 1). In the present study, we selected the putative toxin/immunity pair PAKAF_00796/PAKAF_00797 (PA4163/PA4164) for further characterization, and we refer to it as *tse8-tsi8* (type <u>s</u>ix <u>e</u>xported 8-type <u>s</u>ix immunity 8) in all subsequent sections.

## Tse8-Tsi8 is a novel toxin-immunity pair

To assess the toxic role of Tse8, a strain lacking *tse8* as well as the downstream putative immunity gene (*tsi8*) was generated in a PAKΔ*retS* background, yielding PAKΔ*retS*Δ*tsei8*. In this mutant, expression of *tse8* from pMMB67HE with and without a C-terminal HA tag affected growth (Fig. 2a). Furthermore, in a competition assay this mutant strain carrying a *lacZ* reporter gene (recipient PAKΔ*retS*Δ*tsei8::lacZ*) was outcompeted only by donor strains having an active H1-T6SS, *i.e.* PAKΔ*retS* or PAKΔ*retS*ΔH2ΔH3 (Fig. 2b). The observed killing of the receiver strain was further demonstrated to be Tse8 dependent in competition assays with an attacker lacking Tse8 (Extended Data Fig. 2a). The PAKΔ*retS* strain lacking either *tsei8* or *tse8* could be complemented in a competition assay by expression of *tsei8* from pBBR-MCS5 or *tse8* from pBBR-MCS4 (Extended Data Fig. 2b, c).

**Figure 2.**
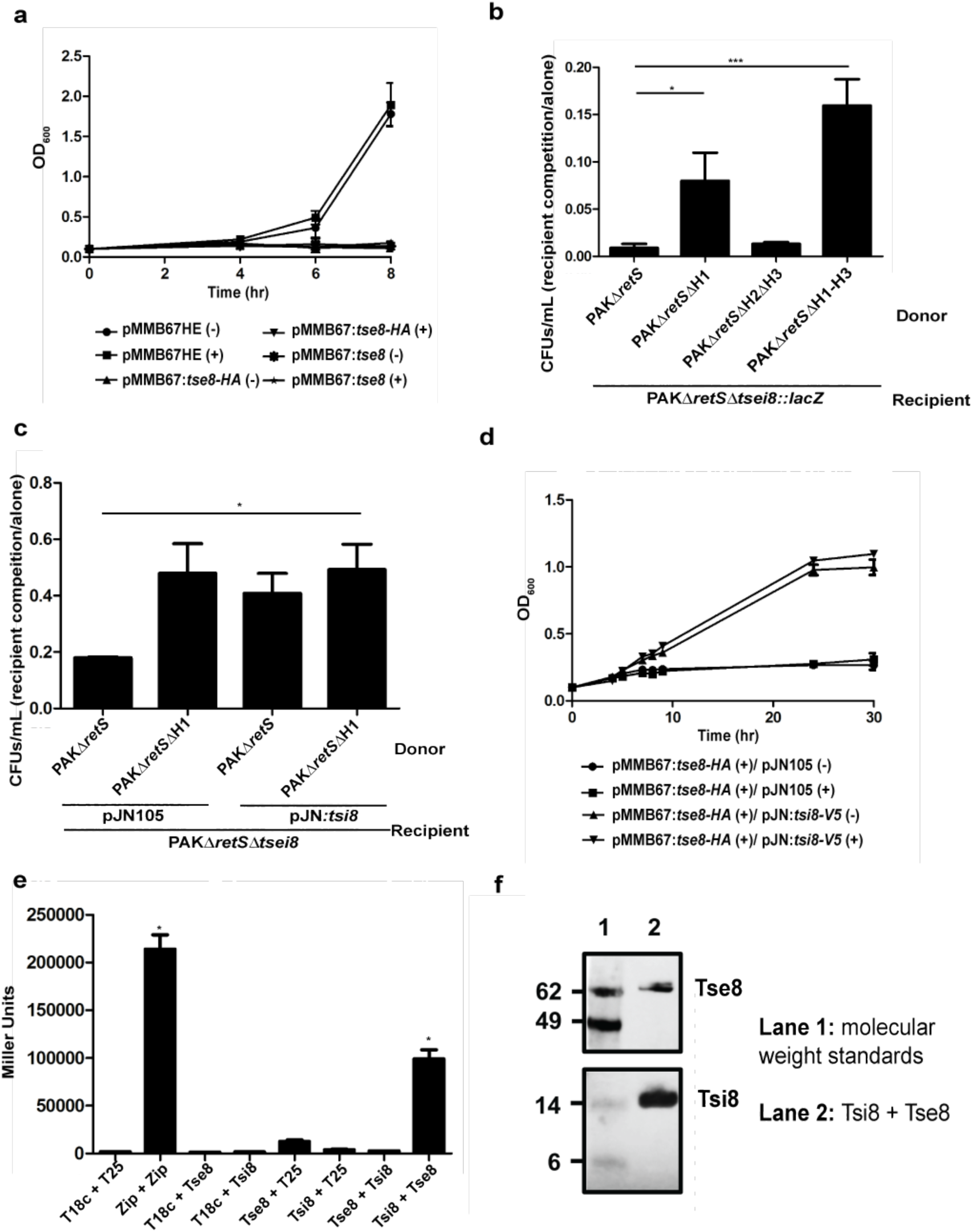
Tse8-Tsi8 is a novel H1-T6SS toxin-immunity pair. **a-b**, Expression of Tse8 (either HA tagged or untagged) in PAKΔ*retS*Δ*tsei8* is toxic when expressed *in trans* from pMMB67HE ((-) no induction; (+) with induction) (**a**) or when delivered by the H1-T6SS into a recipient strain lacking *tsi8* (**b**). **c-d,** Tsi8 can rescue Tse8 toxicity in competition assays with donors PAKΔ*retS* or PAKΔ*retS*Δ*H1* and recipient PAKΔ*retS*Δ*tsei8* expressing either pJN105 or pJN:*tsi8* **(c)** and in growth assays with PAKΔ*retS*Δ*tsei8* expressing pMMB:*tse8* or pJN:*tsi8* **(d)**. **e,** Bacterial-Two-Hybrid (BTH) assays were used to quantify the level of interaction between Tse8 and Tsi8 with β-galactosidase activity assays performed on the cell lysates of each interaction pair. **f,** Tse8-HA-Strep interacts directly and specifically with Tsi8-His. The proteins were mixed and added to His-Tag Dynabeads, using Tsi8-His as a bait. The interaction is Tsi8-His specific (see Extended Data Fig. 3).

The toxicity associated with the H1-T6SS-dependent delivery of Tse8 into a sensitive receiver strain could be rescued by expressing the *tsi8* immunity gene from pJN105 in a competition assay (Fig. 2c) or in a growth assay (Fig. 2d), further confirming the protective role of Tsi8. In several cases, T6SS immunity proteins have been shown to directly interact with their cognate toxins^15, 21, 22^. Here, bacterial-two-hybrid (BTH) assays demonstrate that indeed Tse8 interacts strongly with Tsi8 (Fig. 2e). In addition, pull-down experiments using Tsi8-His as a bait, show direct interaction of the two proteins (Fig. 2f); this interaction is specific to Tsi8 as minimal amounts of Tse8-HA-Strep elute from the pull-down beads in the absence of Tsi8 or in the presence of the non-specific binding control (CcmE-His) (Extended Data Fig. 3).

T6SS toxin delivery frequently relies on a direct interaction between the toxin and components of the T6SS spike^9, 12^. BTH assays (Fig. 3a), as well as far-western dot blots revealed that Tse8 interacts strongly with VgrG1a (Fig. 3b). While the interaction of Tse8 with VgrG1c was significant in the BTH assay (Fig. 3a), no interaction above the non-specific binding control (CcmE-His) was observed in the far-western dot blots (Fig. 3b). Finally, no interaction between Tse8 and VgrG1b was observed in BTH (Fig. 3a) or far western dot blots (Fig. 3b).

**Figure 3.**
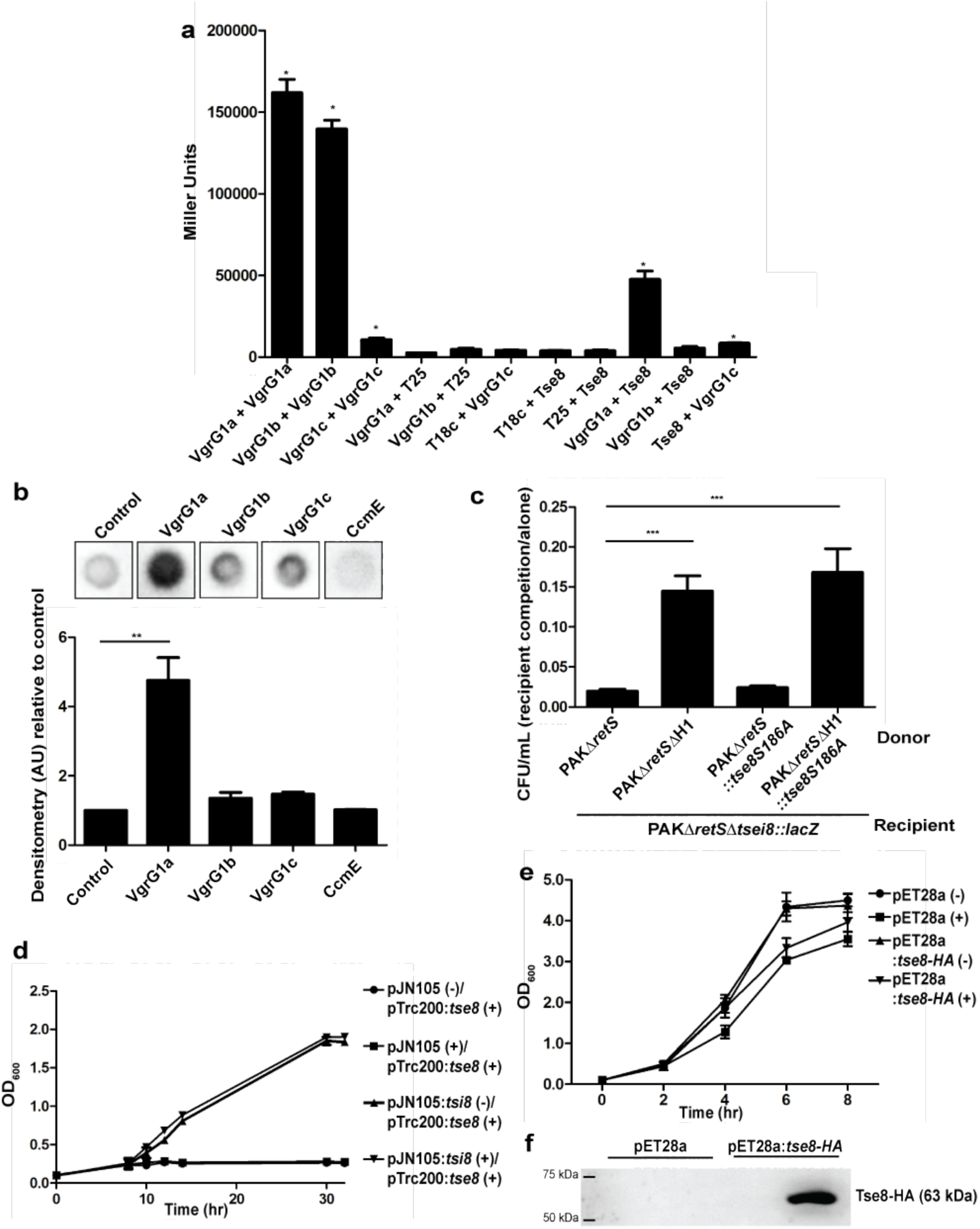
Tse8 interacts with VgrG1a and targets the transamidosome complex. **a**, BTH assays were used to quantify the level of interaction between Tse8 and VgrGs with β-galactosidase activity assays performed on the cell lysates of each interaction pair. **b,** Tse8 interacts with VgrG1a in far western dot blot assays (top panel). Densitometry quantifications of Tse8 interactions with respective partners (bottom panel). CcmE-His is used as a non-specific binding control. **c,** Tse8 toxicity is not dependent on the conserved putative catalytic residue S186. Competition assays were performed with donors PAKΔ*retS,* PAKΔ*retS*ΔH1, PAKΔ*retS::tse8S186A* or PAKΔ*retS*ΔH1::*tse8S186A* and recipient PAKΔ*retS*Δ*tsei8*::*lacZ*. **d**-**f**, Tse8 is only toxic in bacteria which rely on the transamidosome for protein synthesis. Expression of Tse8 in *A. tumefaciens* is toxic but can be rescued by coexpression of Tsi8 ((-) no induction; (+) with induction) (**d**). Expression of Tse8 in *E. coli* is not toxic ((-) no induction; (+) with induction) (**e**), despite Tse8 being expressed (**f**).

Overall, the above results demonstrate that Tse8-Tsi8 is a novel antibacterial toxin-immunity pair associated with the H1-T6SS, and that Tse8 interacts with the VgrG1a tip to facilitate delivery into target cells.

## Tse8 is a predicted amidase family enzyme

Using Phyre2^23^ we found that the closest 3D homologs of Tse8 are the *Stenotrophomonas maltophilia* Peptide amidase (Pam)^24^ (sequence identity 29%), the *Staphylococcus aureus* Gln-tRNA(Gln) transamidosome subunit A (GatA)^25^ (sequence identity 20%), the *P. aeruginosa* Asn-tRNA(Asn) transamidosome subunit A (GatA)^26^ (sequence identity 25%), the *Flavobacterium sp.* 6-aminohexanoate cyclic dimer hydrolase (NylA)^27^ (sequence identity 24%), the *Bradyrhizobium japonicum* malonamidase E2 (MAE2)^28^ (sequence identity 25%), the *Pseudomonas sp.* allophanate hydrolase (AtzF)^29^ (sequence identity 30%), and the *Bacterium csbl00001 Aryl Acylamidase* (AAA)^30^ (sequence identity 22%). Amino acid sequence analysis indicates that Tse8 contains an Amidase Signature (AS) domain (Pfam PF01425) distributed between an N-terminal (residues 25-291) and a C-terminal region (residues 459-544) of its sequence (Extended Data Fig. 4). AS sequences are characterized by a stretch rich in glycine and serine residues, as well as a highly conserved Ser-*cis*Ser-Lys catalytic triad^24, 25, 31–34^. The catalytic Lys is located in the C-terminal end of a conserved β-strand (region 1) (Extended Data Fig. 4), while the *cis*Ser is located at the C-terminus of region 2 (Extended Data Fig. 4). Finally, the nucleophilic Ser residue is located in a highly conserved short loop of region 3. All these AS signature sequence characteristics (underlined by a dashed line in Extended Data Fig. 4) are present in Tse8 and its closest 3D homologues.

Given that Tse8 possesses the conserved catalytic features of amidase family enzymes (Extended Data Fig. 4), we tested whether it has amidase activity. Tse8 was purified and confirmed to be intact (Extended Data Fig. 5). Subsequently, its capacity to hydrolyze carbon-nitrogen bonds was tested on two molecules, epinecidin-1 and glutamine which are substrates for Pam from *S. maltophilia* and GatA of the transamidosome, respectively. The amidase activities of Pam and Tse8 were analyzed by Mass Spectrometry (MS) by monitoring the modifications of epinecidin-1 in the presence and absence of the tested proteins and of the small nucleophile hydroxylamine (Extended Data Fig. 6). While the C-terminus of epinecidin-1 was deaminidated in the presence of Pam (Extended Data Fig. 6b), it remained amidated in the presence of Tse8, suggesting that Tse8 has no amidase activity on this substrate (Extended Data Fig. 6a). The amidase activity of Tse8 was also tested on the GatA substrate glutamine (Extended Data Fig. 7) and no modification was detected by MS (Extended Data Fig. 7b. In addition, whole-cell glutaminase assays were performed and the amidase activity of *E. coli* whole cell lysates expressing GatA or Tse8 on L-glutamine was determined by monitoring the accumulation of NADPH. These experiments demonstrated that while GatA expressed from plasmid pET41a had a significant amount of amidase activity, whole cells expressing Tse8 from the same vector produced a level of NADPH which was not significantly different to the empty vector-carrying control strain (Extended Data Fig. 7c). Overall these data demonstrate that the substrates for Pam and GatA are not substrates for Tse8, suggesting that Tse8 is highly specific or unlikely to utilize amidase activity to elicit toxicity.

To assess whether Tse8 toxicity is mediated through amidase activity *in vivo*, we replaced the *tse8* gene in the chromosome by an allele encoding a putative catalytic site mutant of Tse8 with a Ser186Ala (S186A) substitution. This conserved Ser186 residue (Extended Data Fig. 4) acts as the catalytic nucleophile in homologous amidases, and is necessary for enzymatic function^35^. PAKΔ*retS* and PAKΔ*retS*ΔH1 donor strains encoding either wild-type Tse8 or Tse8S186A were competed against the recipient strain PAKΔ*retS*Δ*tsei8::lacZ*. This showed that there was no difference in the recovered CFUs/mL of the recipient when the attacking strain delivered either wild-type Tse8 or Tse8S186A (Fig. 3c), further suggesting that Tse8 does not utilize amidase activity to elicit toxicity *in vivo*.

## Tse8 elicits toxicity by interacting with the bacterial amidotransferase complex

Since Tse8 toxicity does not appear to depend on it having amidase activity (Fig. 3c), we hypothesized that Tse8 could instead be eliciting toxicity by competing with a functional amidase either within the cell, or within a complex in the cell. Two 3D homologues of Tse8 are the A subunit of the *S. aureus* Gln-tRNA(Gln) transamidosome and the *P. aeruginosa* Asn-tRNA(Asn) transamidosome. Each of these are the A subunit of the transamidosome complex, which are used by bacteria that lack the cognate tRNA synthases for asparagine (Asn) and/or glutamine (Gln)^17^. These bacteria utilize a two-step pathway instead, whereby a non-discriminating tRNA synthase generates a misacetylated aspartate- or glutamate-loaded tRNA which is then transaminated by the heterotrimeric amidotransferase enzyme GatCAB, within the transamidosome complex, to leave asparagine or glutamine correctly loaded onto their cognate tRNA. Given that not all bacteria rely on the transamidosome for protein synthesis, we reasoned that if Tse8 toxicity is directed at this enzymatic complex then expression of Tse8 should only be toxic in bacteria which use the transamidosome. *P. aeruginosa* relies on the transamidosome for Asn-tRNA synthesis^36^ and we see a growth defect when Tse8 is expressed on a plasmid or delivered into a strain lacking Tsi8 (Fig. 2a-d). *Agrobacterium tumefaciens* lacks both Asn-tRNA and Gln-tRNA synthases and generates these cognate tRNAs through the transamidosome (Supplementary Table 2), while *E. coli* possesses both the Asn- and Gln-tRNA synthases and does not have a transamidosome complex (Supplementary Table 2). The effect of Tse8 expression was examined and a growth defect was observed for *A. tumefaciens*, which could be rescued by coexpression of Tsi8 (Fig. 3d), but no growth defect was observed for *E. coli* (Fig. 3e) despite Tse8 expression at high levels from pET28a (Fig. 3f). Taken together these data suggest that Tse8 toxicity depends on the presence of the transamidosome.

A structural homology model of Tse8 was generated based on the solved *S. aureus* GatA 3D structure (PDB: 2F2A). Overlaying the Tse8 homology model with the A subunit of the solved *P. aeruginosa* transamidosome structure (PDB: 4WJ3) (Extended Data Fig. 8a) shows that Tse8 likely shares a high level of structural similarity to the A subunit of the complex. Inspection of the homologous residues within the substrate binding pockets of *Sa*GatA versus *Pa*Tse8 revealed that while the catalytic triad residues are conserved, the substrate binding residues (Tyr309, Arg358 and Asp425 in *Sa*GatA)^24^ are not (Extended Data Fig. 8b), supporting the claim that Tse8 does not have the same substrate as GatA (Extended Data Fig. 7). Given the high level of predicted structural similarity between GatA and Tse8 we hypothesized that Tse8 may be able to interact with the transamidosome and could be eliciting toxicity by altering the functionality of this complex. To investigate this, we performed a pull-down experiment using purified proteins. GatCAB was purified as a complex using a Ni-affinity column through histidine-tagged GatB (His-GatB); GatA and GatC also had tags which were appropriate for their detection by western blot (GatA-V5 and GatC-HA). Tse8 was purified separately through a StrepII tag (Tse8-HA-Strep). GatCAB was pulled down in the presence and absence of a 15-fold molar excess of Tse8 via His-GatB on His-Tag Dynabeads. Tse8 was found to copurify with GatCAB (lane 3, Fig. 4a) and, when present, the amount of pulled GatA decreased accordingly (compare the amounts of GatA in lanes 2 and 3, Fig. 4a); the amounts of pulled GatB and GatC did not decrease in the presence of Tse8. This interaction is specific to GatCAB as minimal amounts of Tse8 elute from the pull-down beads in the absence of the transamidosome or in the presence of the non-specific binding control (CcmE-His) (Extended Data Fig. 3). This is further supported by the fact that Tse8 interacts strongly with GatB and GatC in far Western dot blots, but not with GatA or the binding control CcmE (Fig. 4b).

**Figure 4.**
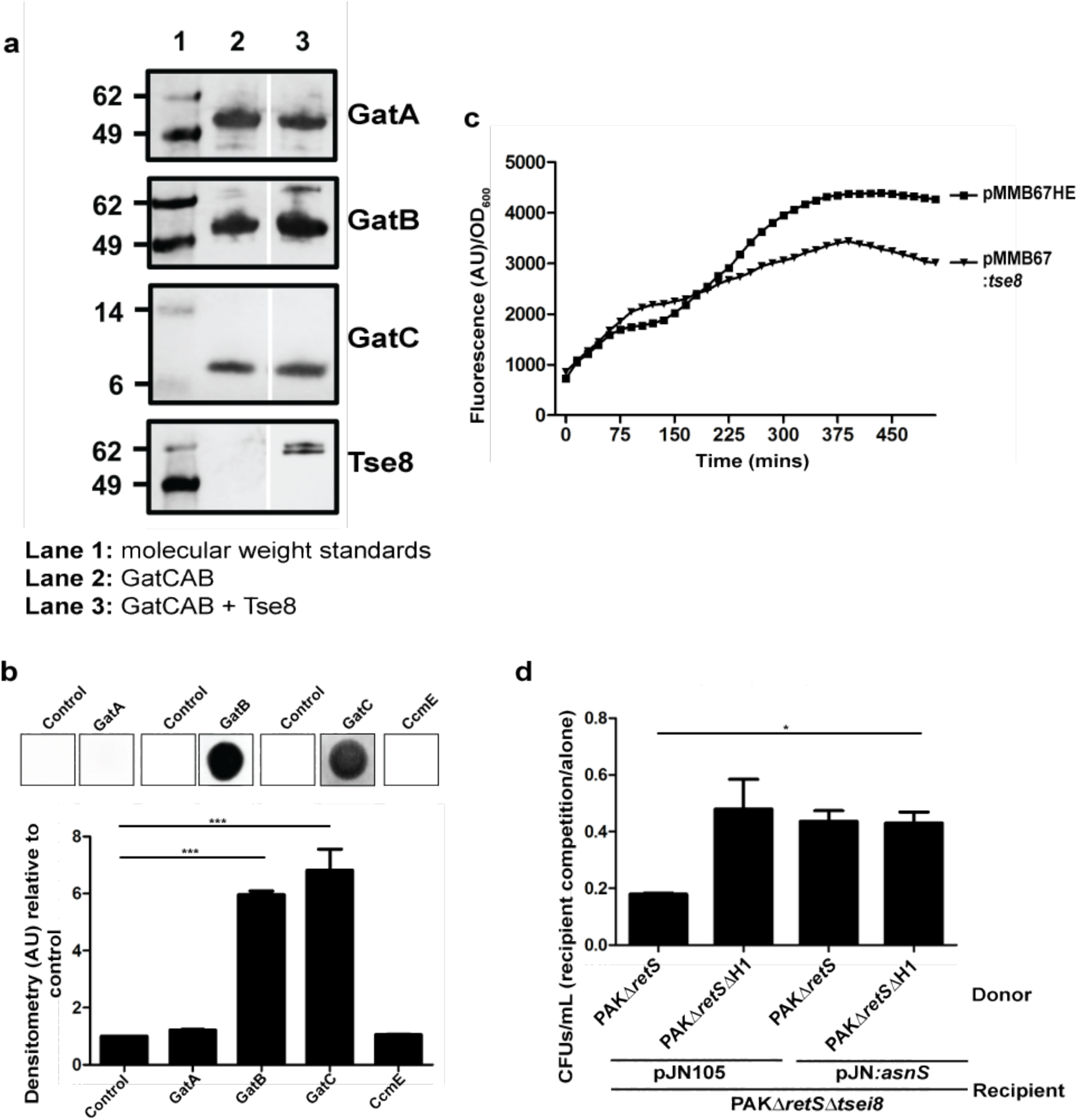
Tse8 interacts with transamidosome components and affects protein synthesis ability. **a**, Tse8-HA-Strep interacts directly and specifically with GatCAB, likely taking the place of GatA in this complex. The proteins were mixed and added to His-Tag Dynabeads, using GatB-His as a bait. Gaps indicate where a lane has been removed. The interaction is GatCAB specific (see Extended Data Fig. 3). **b,** Tse8 interacts with GatB and GatC in far western dot blot assays (top panel). Densitometry quantifications of Tse8 interactions with respective partners (bottom panel). CcmE-His was used as a non-specific binding control. **c,** Representative time course of Gfp levels/OD_600_ within PAKΔ*retS*Δ*tsei8* with an unstable Gfp expressed from the vacant Tn*7* chromosomal site in cells expressing either empty pMMB67HE or pMMB67:Tse8. The data are representative of data obtained in 4 independent experiments. **d,** Asn tRNA synthase (*asnS*) can rescue Tse8 toxicity. Competition assays were performed with donors PAKΔ*retS* or PAKΔ*retS*Δ*H1* and recipient PAKΔ*retS*Δ*tsei8* expressing either pJN105 or pJN:*asnS*.

The structure of the *P. aeruginosa* GatCAB transamidosome reveals it to be a symmetric complex comprising ND-AspRS, GatCAB, and tRNA^Asn^ in a 2:2:2 stoichiometry^26^ (as represented in Extended Data Fig. 8a). Given that Tse8 interacts with the GatCAB complex (Fig. 4a) and specifically with GatB and GatC, but not GatA (Fig. 4b), it could be taking the place of at least one monomer of GatA within one dimer of the GatCAB complex. As Tse8 does not act on the same substrate as GatA *in vitro* (Extended Data Fig. 7), the presence of Tse8 within the complex would reduce the production of Asn-tRNA^Asn^ available for use in protein synthesis. To investigate the effect of Tse8 expression on protein synthesis *in vivo* we utilized an unstable Gfp variant (Gfp-AGA) which is expressed from the Tn*7* site of the *P. aeruginosa* chromosome^37^ in a Tse8-sensitive strain (PAKΔ*retS*Δ*tsei8*) that also expressed Tse8 or the empty pMMB67HE vector. This showed that the levels of Gfp were 30 % lower in the strain expressing Tse8 compared to the empty vector control (Fig. 4c), suggesting that this strain is less able produce the unstable Gfp variant, and thus less cellular protein in general. Given this, we hypothesized that if we were able to override the need for the transamidosome by providing the bacterium with the tRNA synthase it lacked, we would be able to rescue the observed growth defect when Tse8 is either expressed from a plasmid (Fig. 2a,d) or delivered by an attacker (Fig. 2b,c). *P. aeruginosa* only lacks the asparagine tRNA synthase^36^ (and Supplementary Table 2), thus in this case Tse8 toxicity should be rescued by simply providing the cell with this tRNA synthase. To investigate this the Asn-tRNA synthase (*asnS*) from *E. coli* was expressed in PAKΔ*retS*Δ*tsei8* from pJN105, and the strain competed against PAKΔ*retS* and PAKΔ*retS*Δ*H1*. This revealed that expression of AsnS was able to rescue Tse8 toxicity (Fig. 4d) to the same extent as expression of the cognate immunity protein, Tsi8 (Fig. 2c).

## Discussion

In the current study we demonstrate that our global genomic approach can be used to identify T6SS toxin-immunity pairs associated with the H1-T6SS of *P. aeruginosa*. Our approach not only confirmed previously characterized *P. aeruginosa* T6SS toxin-immunity pairs, but also revealed several putative novel toxin-immunity pairs, including Tse8-Tsi8, which would probably not have been found using targeted approaches or bioinformatics. Characterization of the Tse8-Tsi8 pair, revealed that Tsi8 is the cognate immunity protein for the Tse8 toxin, and that Tse8 interacts with VgrG1a, hence it is likely delivered into target cells *via* the VgrG1a-tip complex.

Tse8 was also found to interact with GatCAB of the bacterial transamidosome complex, which is required for protein synthesis in certain bacteria that lack one or both of the asparagine or glutamine tRNA synthases^17^. Our far Western dot blots (Fig. 4b) and pull-down data (Fig. 4a) demonstrate that Tse8 interacts specifically with the GatB and GatC components of the amidotransferase complex, with this interaction likely being mediated by replacement of GatA by Tse8. Nonetheless, the large molar excess required for the pull-down experiments (Fig. 4a) suggests that *in vivo* Tse8 is more likely to interact with transamidosome components as the GatCAB complex assembles *de novo*, rather than displace GatA on already formed transamidosome complexes. Based on this, we propose that Tse8 combines with GatB and GatC as a non-cognate component of the transamidosome complex and that in bacteria where the transamidosome is essential (*i.e*. in bacteria lacking one or both of the Asn- or Gln-tRNA synthases), the formation of such a complex results in reduced fitness due to decreased levels of protein synthesis. In agreement with this, Tse8 toxicity can be rescued if the transamidosome function is bypassed upon provision of the transamidosome-independent tRNA-synthase lacked by the bacterium (*e.g.* AsnS for *P. aeruginosa* (Fig. 4d)).Thus, Tse8 is the first identified T6SS toxin to target a cellular component involved in protein synthesis.

Future work will focus on further characterization of the specifics of the Tse8-GatCAB interaction. This could point to ways of inhibiting the transamidosome and could provide a basis for the development of antibacterial agents against this target. Moreover, investigation of the other putative toxins detected in this study could also open new therapeutic avenues; elucidation of the substrates of these putative toxins could offer insights into pathways that are naturally validated antibacterial targets against *P. aeruginosa*. In looking beyond the T6SS of *P. aeruginosa*, there is a large number of Gram-negative bacteria which infect human and animal hosts, or are plant pathogens or plant-associated organisms and possess at least one, if not multiple T6SSs clusters ^38–42^. Furthermore, in several cases it has been demonstrated that distinct T6SS machines deliver a specific subset of toxins into target cells, often under certain conditions^9, 12, 16^, suggesting that toxins are not only bacterial specific, but potentially even niche specific. Given this diversity, we predict that our TraDIS approach could be useful for drastically expanding the repertoire of known T6SS toxins across a range of bacteria and ecologically or clinically relevant growth environments.

## Methods

### Bacterial strains, plasmids and growth conditions

Bacterial strains and plasmids used in this study are reported in Extended Table 1. *P. aeruginosa* PAK was used for TraDIS library generation and subsequent assays using mutant strains generated by allelic exchange mutagenesis as described previously^43, 44^. *P. aeruginosa* strains were grown in tryptone soy broth (TSB), Lysogeny Broth (LB) or M9 or MOPs minimal media (with indicated supplements) with antibiotics as appropriate (streptomycin 2000 μg/mL, carbenicillin 100 μg/mL, gentamicin 50 μg/mL) at 37 °C with agitation. *E. coli* strains DH5α, SM10, CC118λ*pir* and BL21(DE3) were used for cloning, conjugation and protein expression steps. *E. coli* cells were grown in TSB, LB, Terrific Broth or M9 minimal media (with indicated supplements) (streptomycin 50 μg/mL, ampicillin 100 μg/mL, kanamycin 50 μg/mL) at 37 °C with agitation. *A. tumefaciens* C58 was grown in LB or M9 minimal media (with indicated supplements) with antibiotics as appropriate (gentamicin 50 μg/mL, spectinomycin 100 μg/mL) at 30 °C with agitation.

### DNA manipulation

DNA isolation was performed using the PureLink Genomic DNA mini kit (Life Technologies) except for TraDIS library genomic DNA isolation (see below). Isolation of plasmid DNA was carried out using the QIAprep spin miniprep kit (Qiagen). Primers (Sigma) used are shown in Extended Table 2. DNA fragments were amplified with either KOD Hot Start DNA Polymerase (Novagen) or standard Taq polymerase (NEB) as described by the manufacturer, with the inclusion of Betaine (Sigma) or DMSO (Sigma). Restriction endonucleases (Roche) were used according to the manufacturer’s specifications. DNA sequencing was performed by GATC Biotech.

### TraDIS library generation

A highly saturated transposon mutant library was generated in *P. aeruginosa* PAKΔ*retS* or PAKΔ*retS*ΔH1 strains by large scale conjugation with an *E. coli* SM10 [pBT20] donor which allowed for random insertion of a mariner transposon throughout the genome and conferred gentamicin resistance in the recipient PAK strain. See Supplementary Information for detailed protocol on the generation of the transposon mutant libraries.

### TraDIS library assay

Glycerol stocks of harvested PAKΔ*retS* or PAKΔ*retS*ΔH1 TraDIS libraries were combined at normalized cell density for each separate replicate and spread onto large square (225 mm) VBM agar plates supplemented with gentamicin (60 µg/mL) and incubated for 16 hrs at 37 °C. Cells were then harvested into 5 mL LB and pelleted by centrifugation (10,000 *g*, 15 mins, 4 °C). Cell pellets were resuspended in 1.4 mL LB and 1 mL was taken for subsequent genomic DNA extraction (see below).

### TraDIS library genomic DNA extractions

Genomic DNA from the harvested pooled library pellets were resuspended in 1.2 mL lysis solution (10 mM Tris-HCl, 400 mM NaCl and 2 mM Na_2_EDTA, supplemented with Proteinase K in storage buffer (50 mM Tris-HCl, 50% (v/v) glycerol, 100 mM NaCl, 0.1 mM EDTA, 10mM CaCl_2_, 0.1% (v/v) Triton X-100 and 1 mM DTT) to a concentration of 166 µg/mL. Cell lysis was achieved by incubation at 65 °C for 1 hr, with occasional vortexing. The samples were then cooled to room temperature and RNA removed by addition of RNase A (5 µg/mL) and incubation at 37 °C for 80 mins. Samples were then placed on ice for 5 mins. Each lysate was then split into 2 eppendorf tubes at ∼600 µL per tube, and 500 µL NaCl (5 M) were added to each tube. Cell debris were removed by centrifugation (10,000 *g*, 10 mins, 4 °C) and 500 µL from each tube was added to 2 volumes of isopropanol to precipitate the DNA. DNA was then collected by centrifugation (10,000 *g*, 10 mins, 4 °C), and DNA pellets were washed twice in 70% (v/v) ethanol. The fully dried DNA pellet was finally resuspended in Tris-EDTA buffer.

### PAK reference genome

The PAK genome under the NCBI number accession number LR657304, also listed in the European Nucleotide Archive (ENA) under accession number ERS195106, was used. See details in Cain *et. al.* (2019) *Microbiology Resource Announcement* (submitted 24^th^ July, 2019).

### Generation of TraDIS sequencing libraries, sequencing and downstream analysis

TraDIS sequencing was performed using the method described previously^19^, with some minor modifications for this study. See Supplementary Information for full details.

### Bacterial growth assays

Growth assays were performed as follows. For Fig. 2a, overnight cultures of PAKΔ*retS*Δ*tsei8* were diluted down to OD_600_ = 0.1 in M9 minimal media (with supplements MgSO_4_ (2 mM), CaCl_2_ (0.1 mM), glucose (0.4% (w/v)) and FeSO_4_.7H_2_O (0.01 mM)), and grown shaking at 37 °C. Expression of Tse8 was induced with IPTG (1 mM) at 4 hrs. For Fig 2d, PAKΔ*retS*Δ*tsei8* cells carrying both pJN105 and pMMB67HE plasmids (+/- Tsi8/Tse8) were grown in MOPS minimal media (MOPS (40mM, pH 7.5), Tricine (4 mM, pH 7.5), NH_4_CL (9.52 mM), CaCL_2_ (0.5 uM), MgCl_2_.7H_2_0 (0.52 mM), NaCl (50 mM), FeSO_4_.7H_2_O 20 mM (0.01 mM), K_2_HPO_4_ (1.32 mM) supplemented with 1x micronutrient mix (100x: Ammonium molybdate tetrahydrate (3 uM), Boric acid (400 uM), Cobalt chloride (30 uM), Cupric sulphate (10 uM), Manganese chloride (80 uM), Zinc sulphate (10 uM) and Nickel chloride hexahydrate (0.1% (w/v/)) and glucose (0.4% (w/v)) and L-Glutamine (0.05% (w/v)) without antibiotics shaking at 37 °C. Expression of Tse8 was induced with IPTG (1 mM) and Tsi8 with arabinose (0.2% (w/v)) at 5 hrs. For Fig. 3d, overnight cultures of *A. tumefaciens* with pTrc200/pJN105 plasmids (+/- Tse8/Tsi8) were diluted down to OD_600_ = 0.1 in MOPs media as above without antibiotics and grown shaking at 30 °C. Expression of Tse8 was induced with IPTG (1 mM) and Tsi8 with arabinose (0.2% (w/v)) at 8 hrs. For Fig. 3e, overnight cultures of *E. coli* were diluted down to OD_600_ = 0.1 in M9 minimal media (with supplements MgSO_4_ (2 mM), CaCl_2_ (0.1 mM), FeSO_4_.7H_2_O (0.01 mM) and glucose (0.4% (w/v)) and grown shaking at 37 °C. Tse8 expression was induced by addition of IPTG (1 mM) after 2 hrs.

### T6SS competition assays

T6SS competition assays were performed as described previously^45^ with modifications as indicated. Briefly, overnight cultures of donor and recipient bacteria alone or in a 1:1 ratio were combined and spot plated on LB agar plates for 5 hrs at 37 °C and recovered in serial dilution on LB agar plates supplemented with Xgal (5-bromo-4-chloro-3-indolyl-β-D-galactopyranoside) (100 µg/mL) to differentiate recipient (PAKΔ*retS*Δ*tsei8*::*lacZ* seen as blue) from donor (white). For recovery of competition assays between donor and recipient PAKΔ*retS*Δ*tsei8* [pBBR1-MCS5] and [pBBR1:*tsei8*], the competition assay was plated onto LB agar plates with gentamicin (50 µg/mL) to differentiate donor from recipient (Gm^R^). For recovery of competition assays between donor and recipient PAKΔ*retS*Δ*tsei8* [pBBR1-MCS4] and [pBBR4:*tse8*], the competition assay was plated onto LB agar plates with carbenicillin (50 µg/mL) to differentiate donor from recipient (Carb^R^). In other cases, expression of Tsi8 or AsnS in the recipient strains was induced in the overnight cultures by addition of arabinose (0.2% (w/v)). These overnight cultures of donor and induced recipient alone or in a 1:1 ratio were combined and spot plated onto LB agar supplemented with arabinose (1% (w/v)) for induction of Tsi8-V5 or AsnS-His for 5 hrs, with the competition assay finally being recovered on LB agar plates supplemented with gentamycin (50 µg/mL) and arabinose (1% (w/v)).

### Bacterial Two Hydrid (BTH) assays

Protein-protein interactions were analysed using the BTH system as described previously^46^. Briefly, the DNA region encoding the protein of interest were amplified by PCR and were then cloned into plasmids pKT25 and pUT18C, which each encode for complementary fragments of the adenylate cyclase enzyme, as previously described^46^ resulting in N-terminal fusions of T25/T18 from the adenylate cyclase to the protein of interest. Recombinant pKT25 and pUT18c plasmids were simultaneously used to transform the *E. coli* DHM1 strain, which lacks adenylate cyclase, and transformants were spotted onto Xgal (40 µ/mL) LB agar plates supplemented with IPTG (1 mM), Km (50 µg/mL) and Amp (100 µg/mL). Positive interactants were identified after incubation at 30 °C for 48 hrs. The positive controls used in the study were pUT18C or pKT25 derivatives encoding the leucine zipper from GCN4, which forms a dimer under the assay conditions.

### β-Galactosidase assay

The strength of the interactions in the BTH assays was quantified from the β-galactosidase activity of co-transformants scraped from Xgal plates and measured as described previously; activity was calculated in Miller units^46^.

### Western Blot analysis

SDS-PAGE and western blotting were performed as described previously^9^. Proteins were resolved in 8%, 10%, 12% or 15% gels using the Mini-PROTEAN system (Bio-Rad) and transferred to nitrocellulose membrane (GE Healthcare) by electrophoresis. Membranes were blocked in 5% (w/v) milk (Sigma) before incubation with primary antibodies. Membranes were washed with TBST (0.14 M NaCl, 0.03 M KCl and 0.01 M phosphate buffer plus Tween 20 (0.05% v/v)) before incubation with HRP-conjugated secondary antibodies (Sigma). The resolved proteins on the membrane blots were detected using the Novex ECL HRP Chemiluminescent substrate (Invitrogen) or the Luminata Forte Western HRP substrate (Millipore) using a Las3000 Fuji Imager. For Fig. 3f, samples were taken after 8 hrs of growth and expression of Tse8 was assessed by Western blot as above; detection of Tse8 was performed using α-HA antibody.

### Far-western dot blotting

For Tse8 interactions with VgrG1a, VgrG1b, VgrG1c, GatA, GatB and GatC, purified untagged Tse8 was spotted on nitrocellulose membrane (3 mg/ml) and dried at room temperature. Membranes were blocked with TBST with 5% (w/v) milk or 2.5% (w/v) bovine serum albumin for 7 hrs at room temperature. *E. coli* overexpressing VgrG1a-V5, VgrG1b-V5, VgrG1c-V5 (equivalent 150 OD_600_ units), GatA-V5, His-GatB or GatC-HA (equivalent 200 OD_600_ units) were pelleted and then resuspended in 10 mL 100 mM NaCl, 20 mM Tris, 10% (w/v) glycerol, 2% (w/v) milk powder and 0.1% (v/v) Tween-20 (Tween-20 was added after sonication) (pH 7.6) and sonicated. 10 mL of the crude lysates were applied directly to the membranes and incubated overnight at room temperature. The membranes were immunoblotted with anti-V5 (1:5000 Invitrogen), anti-HA (1:5000 Biolegend), or anti-His (1:1000 Sigma) overnight at 4 °C and anti-mouse secondary (1:5000). Quantification of dot blots was performed using the Gel Analyzer plugin in ImageJ^47^. Levels were normalised to the control signal based on 3 independent experiments.

### Pull-down experiments

*E. coli* BL21(DE3) strains expressing simultaneously GatA-V5, GatB-His and GatC-HA were grown in LB at 37°C to an OD_600_ of 0.8 and expression was subsequently induced using 1 mM IPTG (Sigma) for 16 h at 18 °C. *E. coli* BL21(DE3) cells expressing Tse8-HA-Strep were grown in Terrific Broth at 37°C to an OD_600_ of 0.8 and expression was subsequently induced using 1 mM IPTG (Sigma) for 16 h at 30 °C. The same expression strategy was used for *E. coli* BL21(DE3) strains expressing Tsi8-His or CcmE-His except that TSB medium was used. Cell pellets resulting during expression of GatCAB, Tsi8 and CcmE were resuspended in buffer A (50 mM Tris-HCl, 150 mM NaCl, 20 mM imidazole (pH 7.5)) and lysed by sonication after the addition of protease inhibitors (Roche). Cell debris were eliminated by centrifugation (48,000 *g*, 30 mins, 4 °C). Proteins were purified by immobilised metal affinity chromatography using nickel-Sepharose resin (GE Healthcare) equilibrated in buffer A. Proteins were then eluted off the resin with buffer A containing 200 mM instead of 20 mM imidazole. Cell pellets resulting during expression of Tse8 were resuspended in 50 mM Tris-HCl, 150 mM NaCl (pH 7.5) and lysed by sonication after the addition of protease inhibitors (Roche). Tse8-HA-Strep was purified using Strep-Tactin Sepharose (IBA), according to the manufacturer’s specifications.

Pull-down experiments were performed using the above purified protein solutions and His-Tag Isolation & Pull Down Dynabeads (ThermoFischer Scientific). Briefly, the appropriate protein mixtures were generated by mixing 40 μM of the bait protein with 15-fold molar excess of Tse8-HA-Strep; a condition containing solely the same amount of Tse8-HA-Strep was also tested as a negative binding control. The mixtures were incubated at 25 °C with agitation for 1 hr, added to the Dynabeads and processed according to the manufacturer’s specifications. After elution, samples were denatured in 4 x Laemmli buffer and subjected to western blotting as above. Anti-V5 (1:5000 Invitrogen), anti-HA (1:5000 Biolegend), anti-His (1:1000 Sigma), or anti-StrepII (1:3000 IBA) primary antibodies were used along with an anti-mouse secondary (1:5000).

### Whole-cell glutaminase assays

The whole-cell glutaminase activity was measured as described previously^48^ with some modifications as follows. *E. coli* B834 cells containing empty vector, *gatA* or *tse8* in pET41a were grown to OD_600_ ∼ 0.6 when expression was induced by addition of IPTG (0.5 mM) and grown at 18 °C for 16 h. Cells pellets equivalent to 45 OD_600_ units were washed in sodium acetate solution (sodium acetate (100 mM, pH 6), L-glutamine (20 mM)) and resuspended in a final volume of 600 µL sodium acetate solution, and incubated at 37 °C for 30 mins. 20 µL of cells were retained and serially diluted to quantify the CFUs present. The remaining cell volume was then lysed by heating at 99 °C for 3 min. Once cooled to room temperature 100 µL of cell lysate was added to 2 mL of glutamate dehydrogenase solution (sodium acetate (10 mm), NAD^+^ (4 mM), hydroxylamine HCl (400 mM), 30 U of glutamate dehydrogenase (GDH) enzyme (Sigma) in potassium phosphate buffer (100 mM, pH 7.2)) and incubated at 60 °C for 60 mins. 150 µL of the reaction was added to a 96 well clear plate and the relative accumulation of NADPH was calculated using the measured absorbance at 340 nm.

### Expression and purification of Tse8 used for activity measurements

The pET41a::GST-TEV-Tse8 vector coding for *P. aeruginosa* Tse8 was obtained by FastCloning^49^ using pET41a:GST-Tse8 (see Extended Table 1) as template in order to express Tse8. See Supplementary Information for full details of expression and purification of Tse8.

### Tse8 substrate activity assays

Putative Tse8 substrates were selected based on the predicted GatA and PAM homology. Thus, the capacity of Tse8 to hydrolyse carbon-nitrogen bonds was analysed by mass spectrometry (MS) using as putative substrates the free amino acid glutamine and the C-terminally amidated peptide epinecidin-1 (sequence: GFIFHIIKGLFHAGKMIHGLV-NH_2_) (Bachem AG). Glutamine (10 mM) was incubated with 2 μM of freshly-purified Tse8. Reactions were carried out in two different buffers to test the possible effect of pH; one set of reactions was carried out in 10 mM sodium phosphate buffer (pH 7.6) and another set of reactions was carried out in 20 mM Tris-HCl buffer (pH 8.3). For epinecidin-1, 5 μM of freshly-purified Tse8 or the positive control protein Pam (purified as described previously^50^), were incubated with 50 μM of putative substrate in 10 mM sodium phosphate buffer (pH 7.2); control reactions, lacking Tse8 or Pam, were also tested. Reactions were incubated overnight at 30 °C, followed by MS analysis. For full details on the MS analysis see Supplementary Information.

### Bioinformatics analyses

To predict which bacteria possess AsnS and/or GlnS and/or the amidotransferase GatCAB complex *E. coli* AsnS and GlnS protein sequences or *P. aeruginosa* GatA and GatB protein sequences were used to search the National Center for Biotechnology Information (NCBI) collection of non-redundant protein sequences of bacteria and archaea (*non-redundant Microbial proteins*, update: 2017/11/29) using the *pBLAST* search engine. See Supplementary Information file for full details on bioinformatic analysis.

## Data availability statement

PAK genome NCBI number is LR657304 and in the ENA (European Nucleotide Archive) is ERS195106. The resulting sequences of the T6SS TraDIS assays are available from the European Nucleotide archive (ENA) under study accession number ERS577921.

## Statistical analyses

Statistical analysis was performed using GraphPad Prism version 5 and are represented in figures throughout the text as detailed below.

## Figure 2

Figure 2a – Mean OD_600_ ± SEM is plotted over time from 3 independent replicates.

Figure 2b – Mean CFUs/mL ± SEM of recipient cells in competition/alone are represented from 3 independent replicates performed in triplicate. Two-tailed student’s t-test, *** p<0.001; * p<0.05; ns between PAKΔ*retS* and PAKΔ*retS*ΔH2ΔH3 (p = 0.436).

Figure 2c – Mean CFUs/mL ± SEM of recipient cells in competition/alone are represented from 3 independent replicates performed in triplicate. Two-tailed student’s t-test, * p<0.05 for each sample to PAKΔ*retS*; ns between PAKΔ*retS*ΔH1 [pJN105] and PAKΔ*retS* [pJN:*tsi8*] (p=0.598).

Figure 2d – Mean OD_600_ ± SEM is plotted over time from 3 independent replicates.

Figure 2e – Mean ± SEM of three biological replicates performed in triplicate. One-way Anova with Tukey’s multiple comparison post-test, * p<0.05 compared to the Miller units for both T18c:tsi8 + T25 and T18c + T25:tse8.

## Figure 3

Figure 3a – Mean ± SEM of three biological replicates performed in triplicate. One-way Anova with Tukey’s multiple comparison post-test, * p<0.05 compared to the Miller units for each of VgrG1a, VgrG1b, VgrG1c and Tse8 with the respective T18c or T25 partner.

Figure 3b – Densitometry measurements normalized to the control and represented as the Mean ± SEM from 3 independent replicates. Two-tailed student’s t-test, ** p<0.005 compared to control; ns between control and VgrG1b (p=0.169), VgrG1c (p=0.067) and CcmE (p=0.159).

Figure 3c – Mean CFUs/mL ± SEM of recovered recipient are represented from 3 independent replicates performed in triplicate. Two-tailed student’s t-test, *** p<0.001 for PAKΔ*retS* compared to PAKΔ*retS*ΔH1 and PAKΔ*retS*ΔH1::*tse8S186A*; ns between PAKΔ*retS* and PAKΔ*retS::tse8S186A* (p = 0.226).

Figure 3d-e – Mean OD_600_ ± SEM is plotted over time from 3 independent replicates.

## Figure 4

Figure 4b – Densitometry measurements normalized to the control and represented as the Mean ± SEM from 3 independent replicates. Two-tailed student’s t-test, *** p<0.001; ns for control compared to GatA (p=0.077) or to CcmE (p=0.089).

Figure 4d – Mean CFUs/mL ± SEM of recipient cells in competition/alone are represented from represented from 3 independent replicates performed in triplicate. Two-tailed student’s t-test, * p<0.05; ns for PAKΔ*retS*ΔH1 [pJN105] *vs* PAKΔ*retS* [pJN:*asnS*] (p = 0.687) or *vs* PAKΔ*retS*ΔH1 [pJN:*asnS*] (p = 0.631).

## Extended Figure 2

Extended Figure 2a – Mean CFUs/mL ± SEM of recipient cells in competition/alone are represented from represented from 3 independent replicates performed in triplicate. Two-tailed student’s t-test, ** p<0.005 each compared to PAKΔ*retS* donor vs recipient.

Extended Figure 2b – Mean CFUs/mL ± SEM of recipient cells in competition/alone are represented from represented from 4 independent replicates performed in triplicate. Two-tailed student’s t-test, *** p<0.001 compared to PAKΔ*retS* donor vs recipient PAKΔ*retS*Δ*tsei8* [pBBR1-MCS5] compared separately to each other data point; ns between recovered CFUs/mL for recipient PAKΔ*retS*Δ*tsei8* [pMMB-MCS5] *vs* PAKΔ*retS*ΔH1 (p=0.51) and recipient PAKΔ*retS*Δ*tsei8* [pMMB:*tsei8*] *vs* PAKΔ*retS* (p=0.61).

Extended Figure 2c – Mean CFUs/mL ± SEM of recipient cells in competition/alone are represented from represented from 3 independent replicates performed in triplicate. Two-tailed student’s t-test, *** p<0.005 for PAKΔ*retS* [pBBR1-MCS4] vs recipient compared to PAKΔ*retS*Δ*tse8* [pBBR1-MCS4] *vs* recipient; * p<0.05 for PAKΔ*retS* [pBBR1-MCS4] *vs* recipient compared to PAKΔ*retS* [pBBR1:*tse8*] and PAKΔ*retS*Δ*tse8* [pBBR1:*tse8*] *vs* recipient.

## Extended Figure 7

Extended Figure 7c – Mean ± SEM of four biological replicates performed in triplicate. Two-tailed student’s t-test, *** p<0.0001 for empty vector compared to pET41a:*gatA*; ns for empty vector compared to pET41A:*tse8* (p=0.621).

## Supplementary Information

Supplementary information document includes: Materials and methods; Supplementary Table 1 with full list of TraDIS gene hits obtained with log fold change (logFC), p value for the standard and q value and Supplementary Table 2 with a list of relevant bacteria identified with either AsnS/GlnS and/or GatCAB sequence.

## Acknowledgements

The Authors state that they have no conflict of interest. L.M.N. is supported by MRC Grant MR/N023250/1 and a Marie Curie Fellowship (PIIF-GA-2013-625318). A.F. is supported by Medical Research Council (MRC) Grants MR/K001930/1 and MR/N023250/1 and Biotechnology and Biological Sciences Research Council (BBSRC) Grant BB/N002539/1. D.A.I.M. is supported by the MRC Career Development Award MR/M009505/1. D.A.J acknowledges support by the MINECO Contract C16-76941-R. M.A.S.P. is supported by the MINECO under the “Juan de la Cierva Postdoctoral program” (position FJCI-2015-25725). Technical support from the CIC bioGUNE Metabolomics and Proteomics platforms are gratefully acknowledged.

## Author contributions

L.M.N designed the overall experimental plan for the manuscript, performed the majority of the experiments presented and wrote the manuscript; A.K.C performed all TraDIS sequencing and associated bioinformatic analyses; E.M. assisted with protein pull-down assays; M.A.S.P performed protein purification and MS enzymatic assays; D.A.J designed and performed homology modelling, bioinformatic analyses, protein purification and enzymatic assays; D.A.I.M. performed protein purification and pull-down experiments and contributed to the revision of the manuscript; D.A.J performed homology modelling and bioinformatic analyses; G.D and J.P contributed to project management and supported TraDIS sequencing and associated bioinformatic analyses; A.F contributed to project management, designed the overall experimental plan for the manuscript, and contributed to writing the manuscript.

## Extended Tables

**Extended Table 1.**
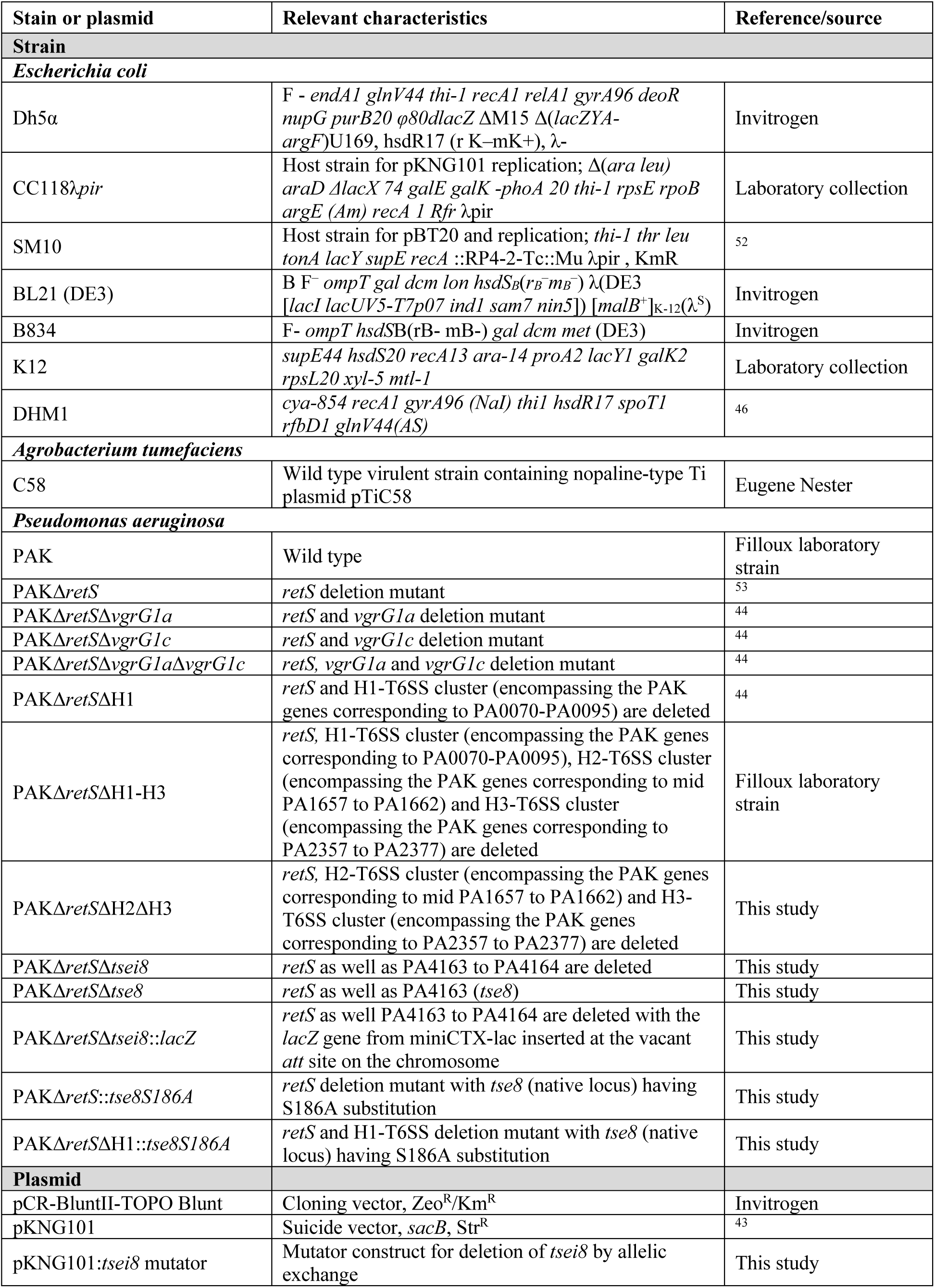
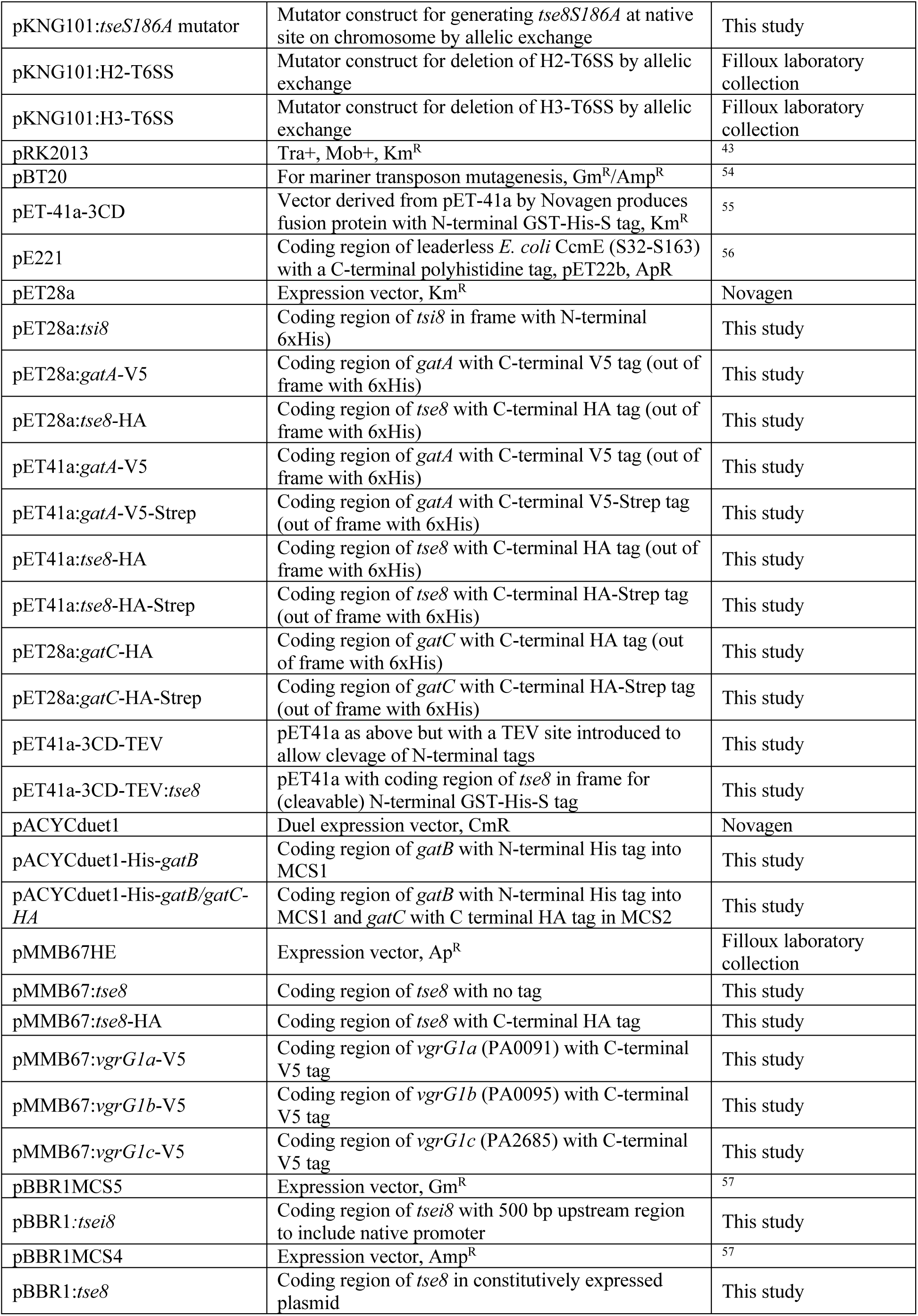
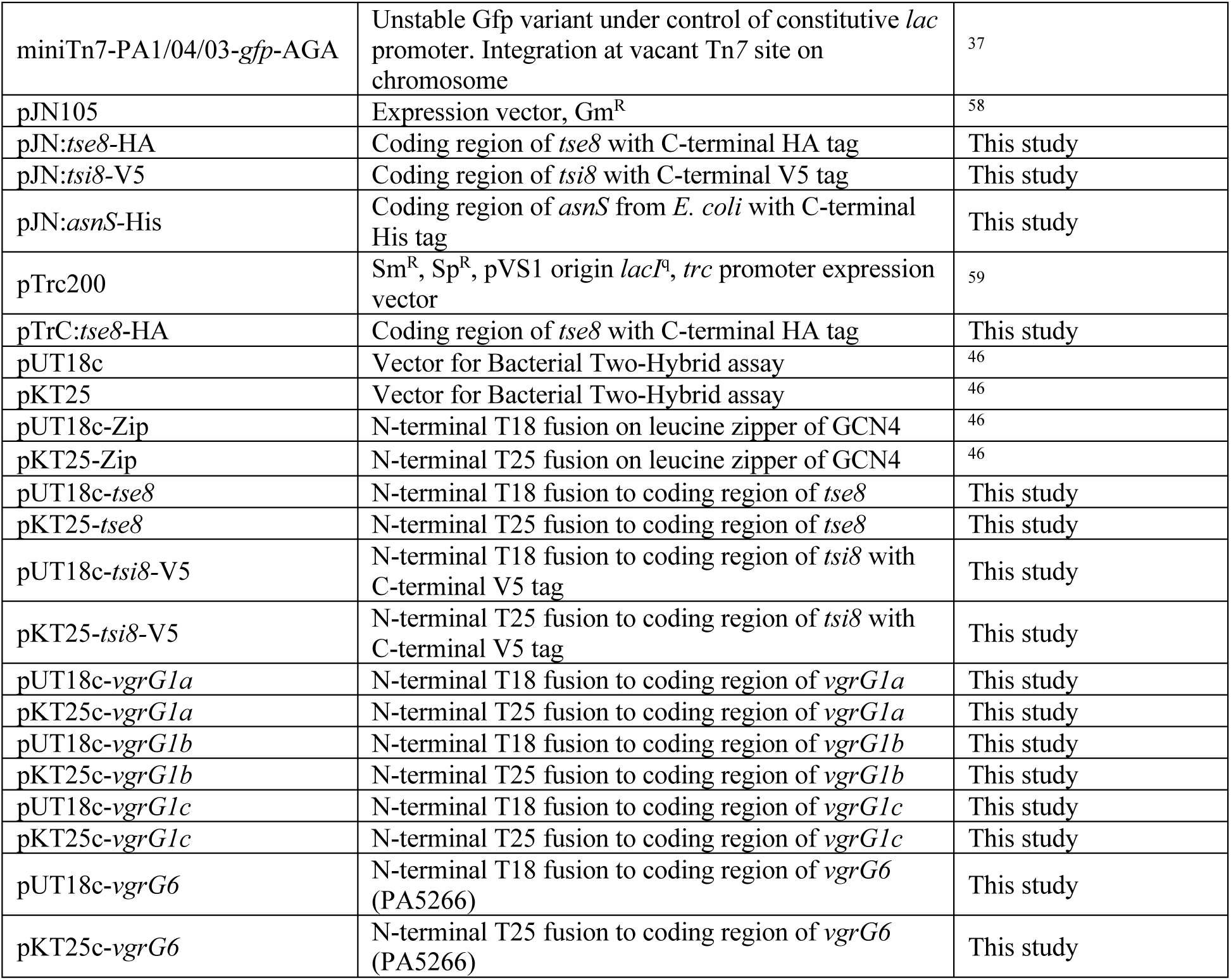
Strains and plasmids used in the current study.

**Extended Table 2.**
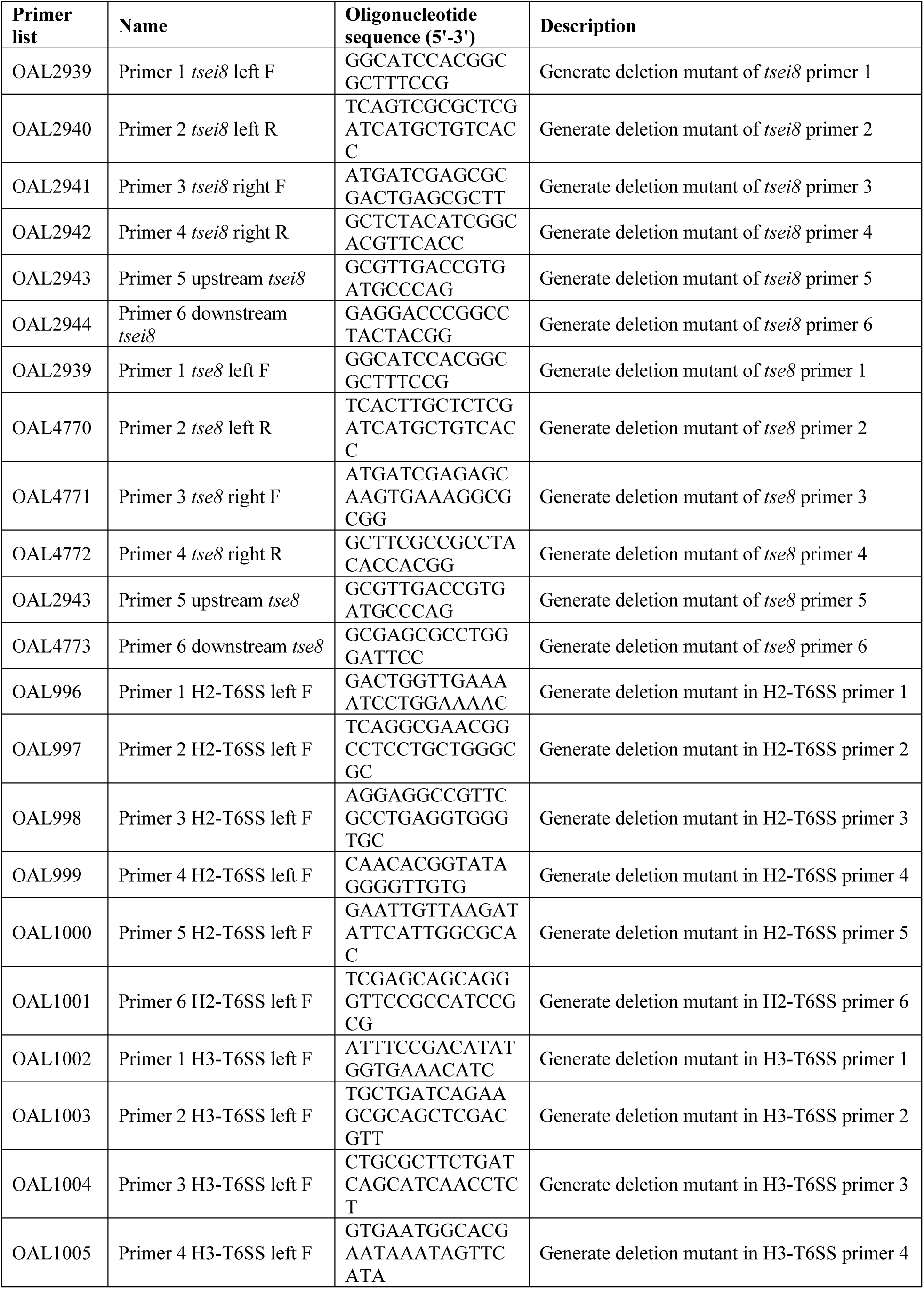
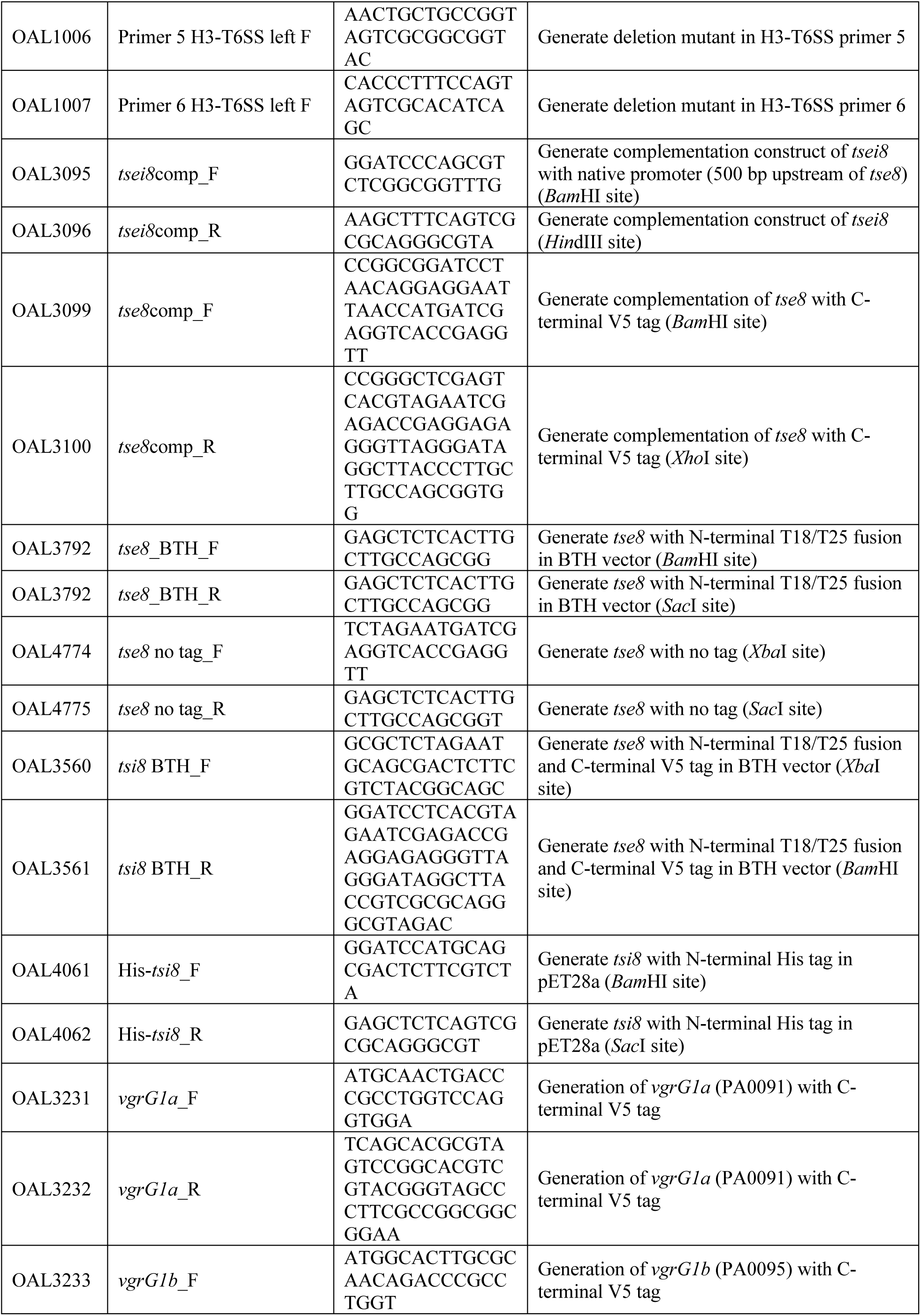
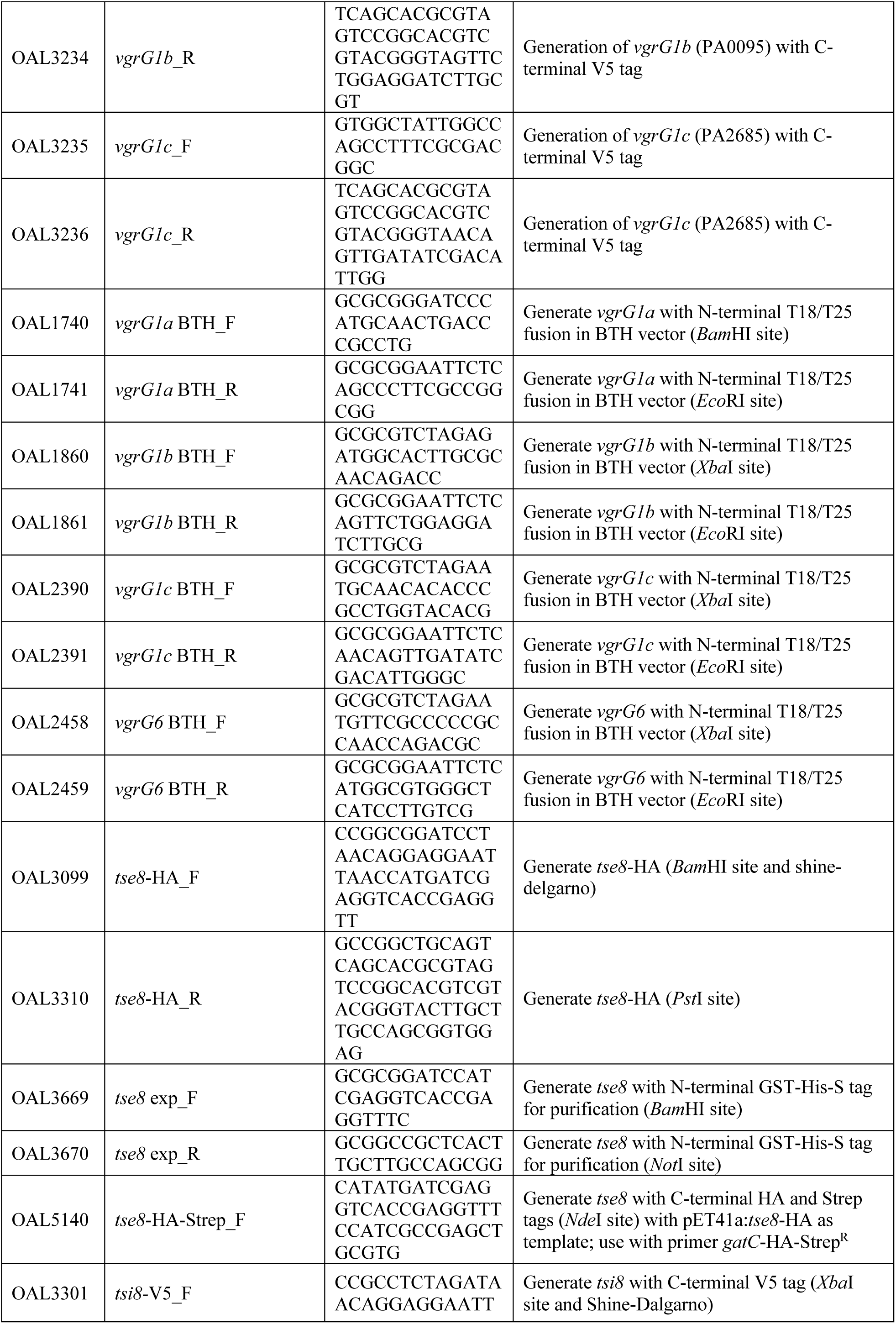
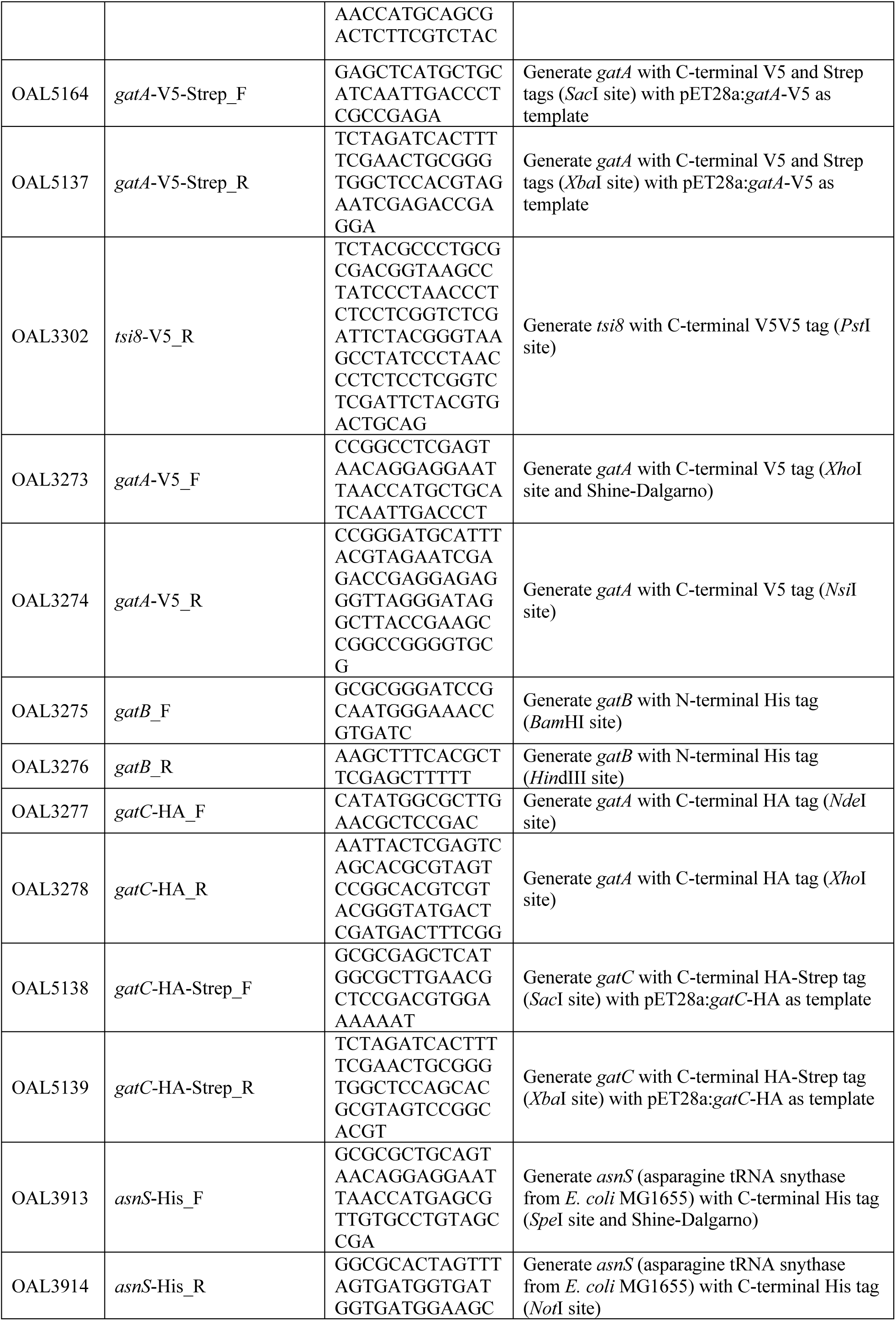
Primers used in the current study.

## Extended Data Figures and Figure Legends

**Extended Data Figure 1.**
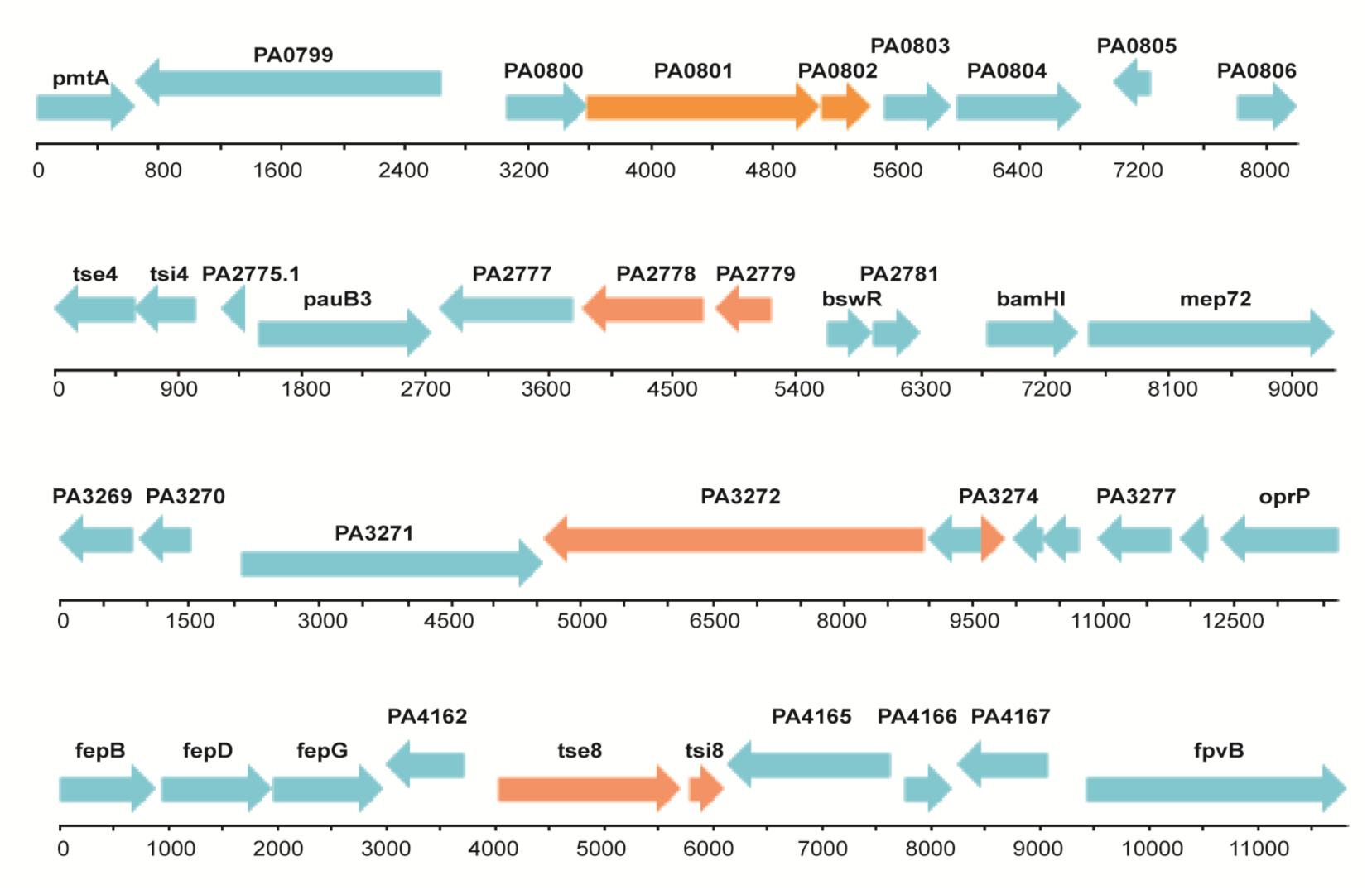
Genomic context of putative novel toxin-immunity pairs identified in TraDIS screen. Putative toxin and immunity pairs from Table 1 are in orange with surrounding genes in blue. Genes corresponding to PAO1 ORF numbers. Base pairs covering the region are marked below each gene sequence.

**Extended Data Figure 2.**
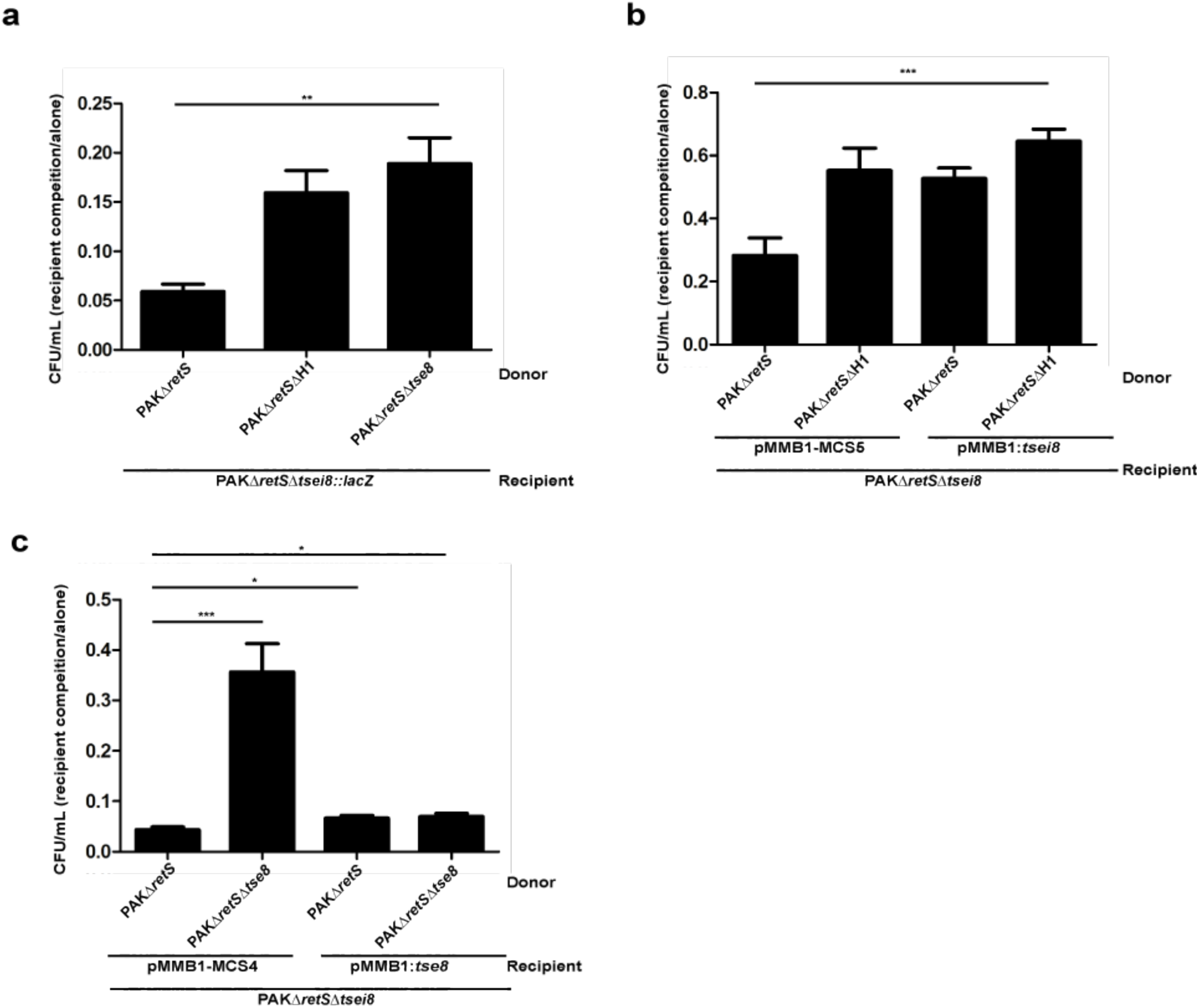
Prey killing is mediated by Tse8 and effects can be complemented by expressing Tse8 or Tsei8 *in trans*. **a**, In the absence of Tse8 (PAKΔ*retS*Δ*tse8*) or the H1-T6SS (PAKΔ*retS*ΔH1) there is no reduction in recovered recipient (PAKΔ*retS*Δ*tsei8*) as occurs when the donor has a fully active T6SS (PAKΔ*retS*). **b-c,** The PAKΔ*retS*Δ*tsei8* **(b)** or PAKΔ*retS*Δ*tse8* **(c)** mutation can be complemented *in trans.* Competition assays were performed with donors PAKΔ*retS* or PAKΔ*retS*Δ*H1* and recipient PAKΔ*retS*Δ*tsei8* with either empty pBBR1MCS5 or the complementation vector pBBR1*:tsei8* **(b)** or recipient PAKΔ*retS*Δ*tsei8* with either empty pBBR1MCS4 or the complementation vector pBBR1*:tse8* **(c)**.

**Extended Data Figure 3.**
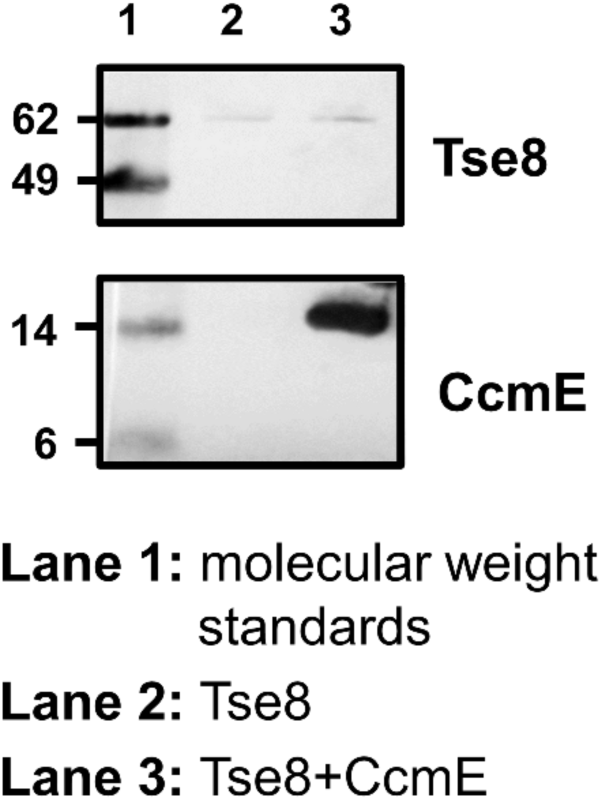
The interaction of Tse8 with Tsi8 or GatCAB is specific. Tse8-HA-Strep does not bind non-specifically to the His-Tag Dynabeads used in the pull-down experiments in Fig. 2f and 4a and does not interact with the non-specific binding control protein (CcmE-His). Tse8-HA-Strep was added to the His-Tag Dynabeads and to the CcmE-His protein solution in the same molar excess as used in the experiments shown in Fig. 2f and 4a. Negligible amounts of Tse8-HA-Strep can be detected in both conditions.

**Extended Data Figure 4.**
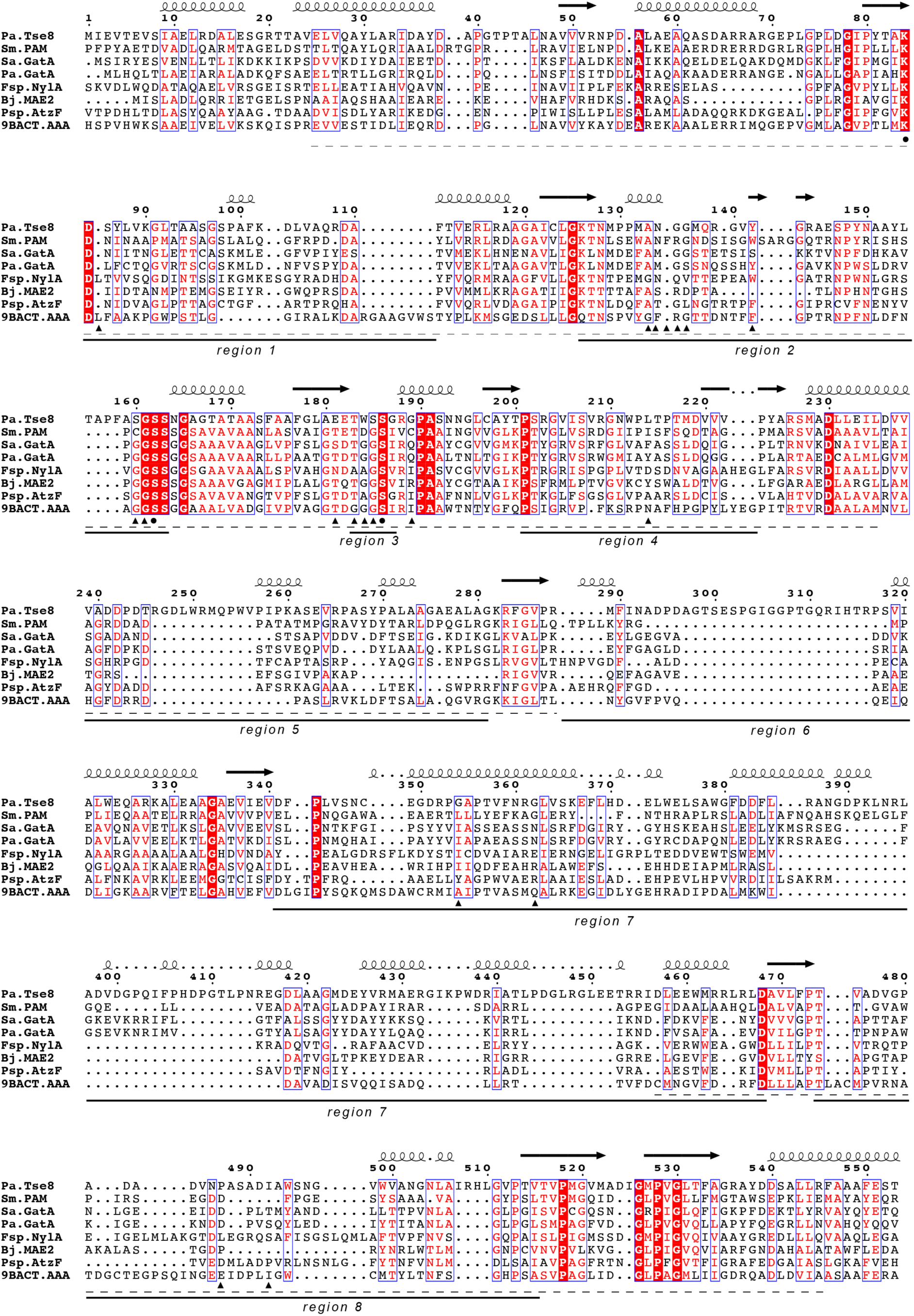
Sequence alignment of Tse8 with predicted homologs of known 3D structure. Amino acid sequences from *P. aeruginosa* Tse8 (*Pa.*Tse8), the *Stenotrophomonas maltophilia* Peptide amidase (*Sm*.Pam), the *Staphylococcus aureus* Gln-tRNA(Gln) amidotransferase subunit A (*Sa*.GatA), the *P. aeruginosa* Asn-tRNA(Asn) transamidosome subunit A (*Pa.*GatA), the *Flavobacterium sp.* 6-aminohexanoate cyclic dimer hydrolase NylA (*Fsp*.NylA), the *Bradyrhizobium japonicum* malonamidase E2 (*Bj*.MAE2), the *Pseudomonas sp.* allophanate hydrolase (*Psp*.AtzF) and the *Bacterium csbl00001* Aryl Acylamidase (*9BACT*.AAA) were aligned. Residues are color-coded depending on the percentage of equivalences; white letter in red background for residue 100 % conserved, red letter in white background for residue with physical-chemical properties conserved. The secondary structure elements found in the 3D structure of *Sm*.PAM are represented above the alignment (black arrows correspond to β-sheets and curly lines to α-helices). The conserved Ser-Ser-Lys catalytic triad is indicated below the alignment by black dots. The AS signature sequence is indicated below the alignment by a dotted line. Regions that protrude out of the core AS domain are numbered below the alignment. Residues found to interact with substrates/substrate analogues, products or inhibitors are indicated with black triangles below the alignment (analysis carried out for crystal structures with the following PDB codes: 1M21 (*Sm*.Pam), 1O9O (*Bj*.MAE2), 4CP8 (*Psp*.AtzF) and 4YJI (*9BACT*.AAA)). Alignment was generated with MUSCLE^60^ and graphical representation with ESPript 3^61^.

**Extended Data Figure 5.**
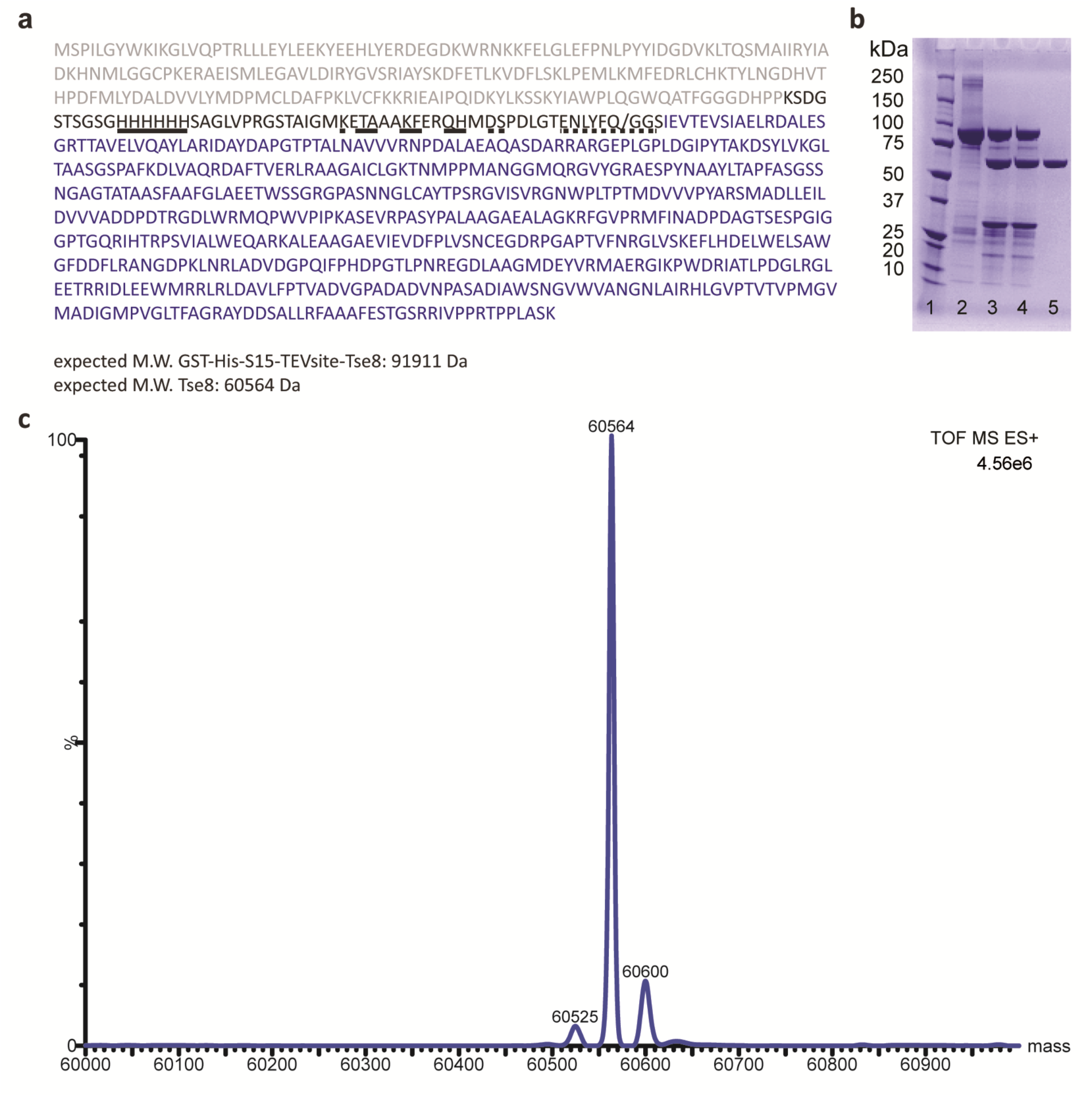
Recombinant production of Tse8. **a**, Amino acid sequence of *Pseudomonas aeruginosa* GST-TEV-Tse8 construct. The recombinant Tse8 construct contains a fused glutathione S-transferase (GST) tag (grey colour), a S15 tag (dashed line), a poly-histidine tag (smooth line) and the optimal Tobacco Etch Virus protease (TEV) cleavage site (ENLYFQG) (dotted line) at the N-terminus of Tse8 (in blue letters). **b**, Sodium dodecyl sulfate polyacrylamide gel electrophoresis (SDS-PAGE) of purified Tse8. Lane 1, molecular weight marker; Lane 2, sample before cleaving with TEV; Lanes 3-4, sample after incubation with TEV; Lane 5, Tse8 without tags (4-20% gel (ExpressPlus™ PAGE Gel, GenScript). **c,** Deconvoluted electrospray ionization mass spectrometry (ESI-MS) chromatogram of purified Tse8 after TEV cleavage (the experimentally determined molecular weight corresponds to the expected molecular weight of 60,564 Da).

**Extended Data Figure 6.**
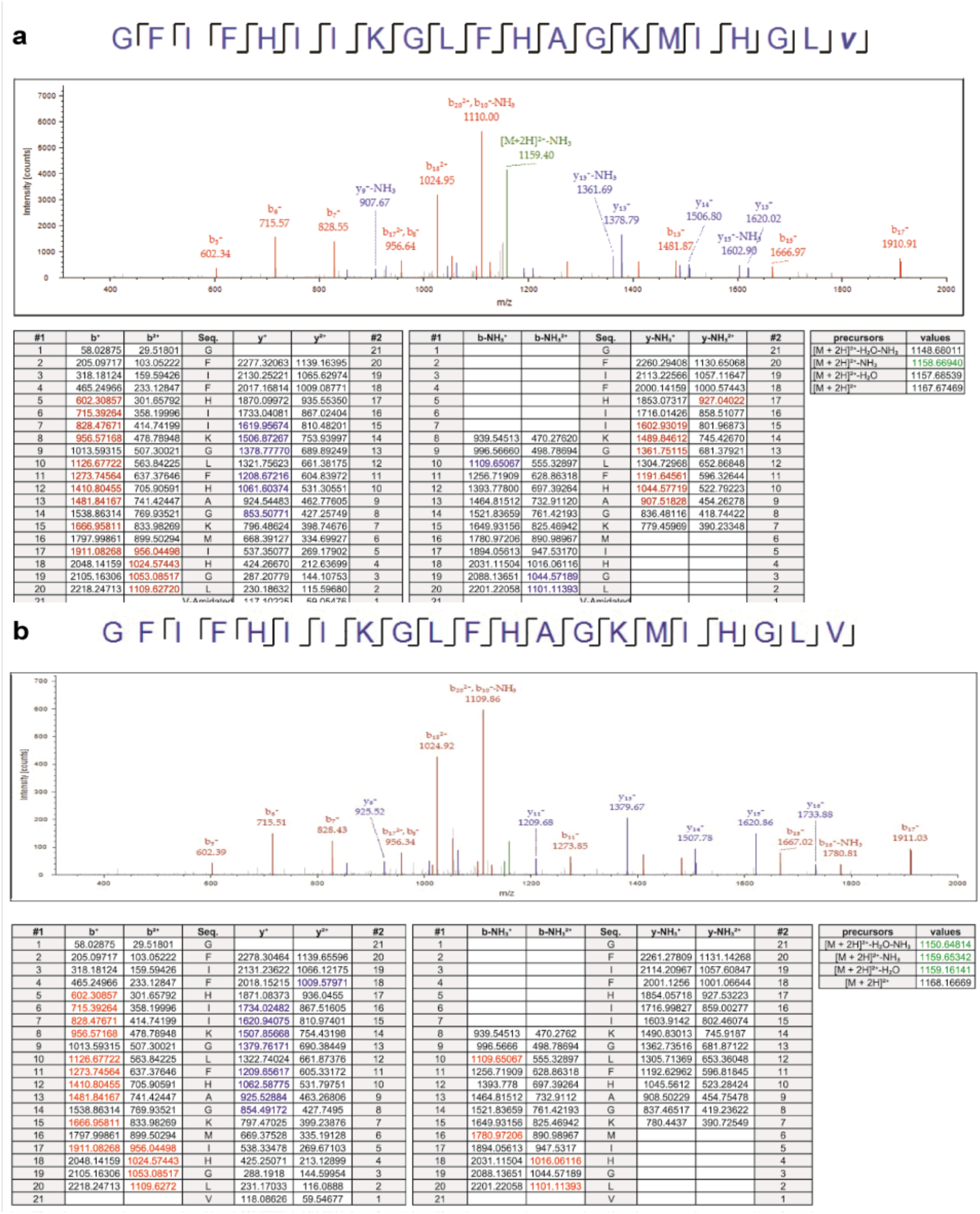
Tse8 is not active on a substrate of the amidase Pam. **a-b,** MS analysis of Tse8 **(a)** or Pam **(b)** enzymatic assay using epinicedin-1 as substrate. The antimicrobial peptide epinecidin-1, as well as sermorelin, have amidated C termini. The latter has previously been used to measure the amidase activity of Pam from *S. maltophilia*^50^. Top panel in both **(a)** and **(b)**: Sequence covered by the fragments obtained after fragmentation of epinecidin-1 in the MS is indicated above the fragmentation spectra for epinecidin-1. Signals corresponding to the amidated **(a)** or deamidated **(b)** form of epinecidin-1 are shown in the spectral plot with red ions belonging to the *b* series of fragments, blue ions to the *y* series, and green ions to parental forms of the peptide. Lower panel in both **(a)** and **(b)**: Correspondence between the observed fragments and their theoretical masses. Tables correspond to unaltered ion series at +1 and +2 charge states (left), ion series after neutral losses at +1 and +2 charge states (ammonia loss, centre) and parental ion masses at +2 charge state with or without diverse neutral losses (right).

**Extended Data Figure 7.**
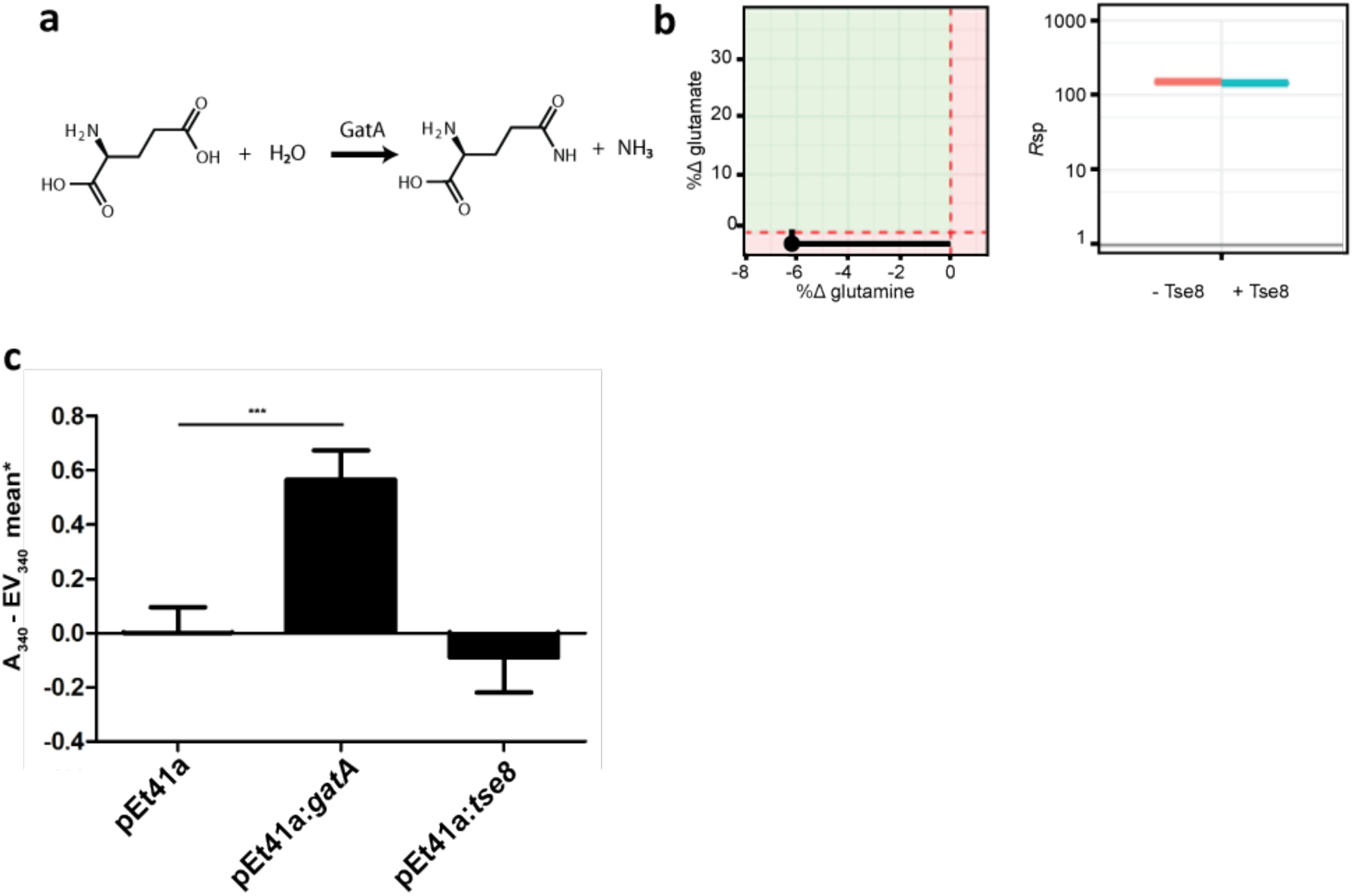
Tse8 is not active on a substrate of GatA. **a**, Amidase reaction catalyzed by GatA and **b,** MS analysis of Tse8 enzymatic assay using glutamine as substrate. Signal of the expected product (glutamate) was also found as a contaminant in the blank and product stock and therefore subtracted from the reaction incubation. The graph on the left shows relative differences (%Δ) of glutamate and glutamine in reaction incubation and blank samples. The green-shaded area indicates the zone in which the observed differences could indicate enzymatic reaction. (%Δ_Product_ = 100 * ([product in incubation] – [product in blank]/ [product in incubation]; %Δ_Substrate_ = 100 * ([substrate in incubation] – [substrate in blank]/ [substrate in incubation]). The graph on the right shows the ratios between substrate and product (*R*_sp_) in the blank (red) and reaction (Tse8) incubation (blue) samples. The product signal (contaminant) was *ca.* 100 times lower than that of the substrate. (*R*_sp_ = (Signal substrate / Signal product)). **c,** Glutaminase assays of lysates of *E. coli* cells expressing GatA, Tse8 or empty vector demonstrate that Tse8 does not have the same substrate (L-glutamine) as GatA as measured by relative NADPH levels/(CFU/mL) (*represented as empty vector (EV) mean subtracted from each mean sample).

**Extended Data Figure 8.**
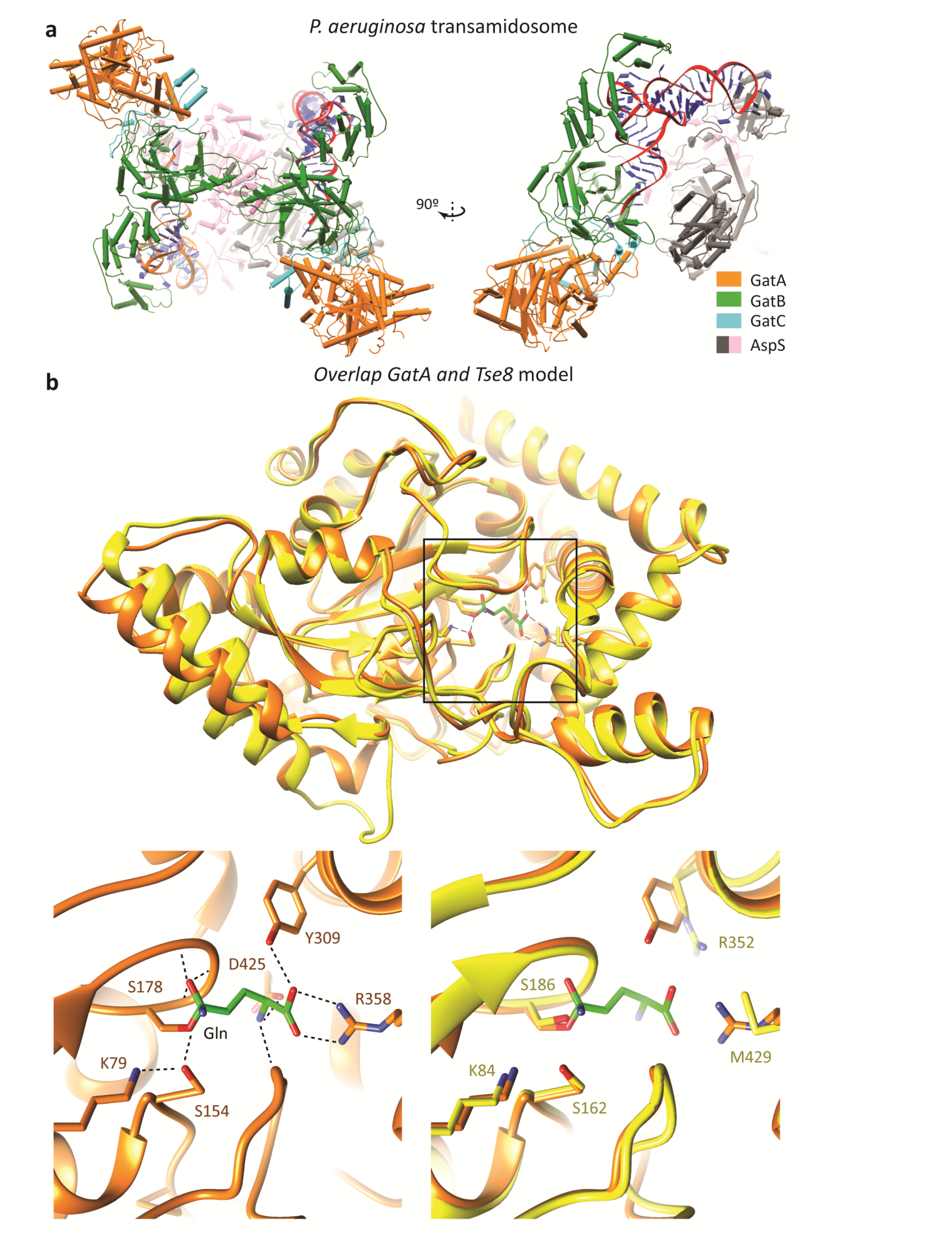
Tse8 is structurally similar to GatA of the transamidosome complex. **a**, Structure of the *P. aeruginosa* GatCAB transamidosome-Asp-tRNA structure (PDB: 4WJ3). **b,** Top panel: Tse8 3D homology model generated using GatA from *S. aureus* (from PDB: 2F2A) as template overlaid with the A subunit of the solved GatCAB transamidosome-AspS-tRNA structure from *P. aeruginosa* (PDB: 4WJ3). The reaction centre with covalently bound glutamine substrate is boxed. Bottom panel: Close-up view of the reaction centre of *S. aureus* GatA (left) with glutamine (green) substrate bound and of a superposition of *S. aureus* GatA and the 3D homology model of *P. aeruginosa* Tse8 (right) showing the predicted conservation of the Ser-*cis*Ser-Lys catalytic triad and predicted divergent substrate binding residues in Tse8 compared to GatA.

**Extended Figure 9.**
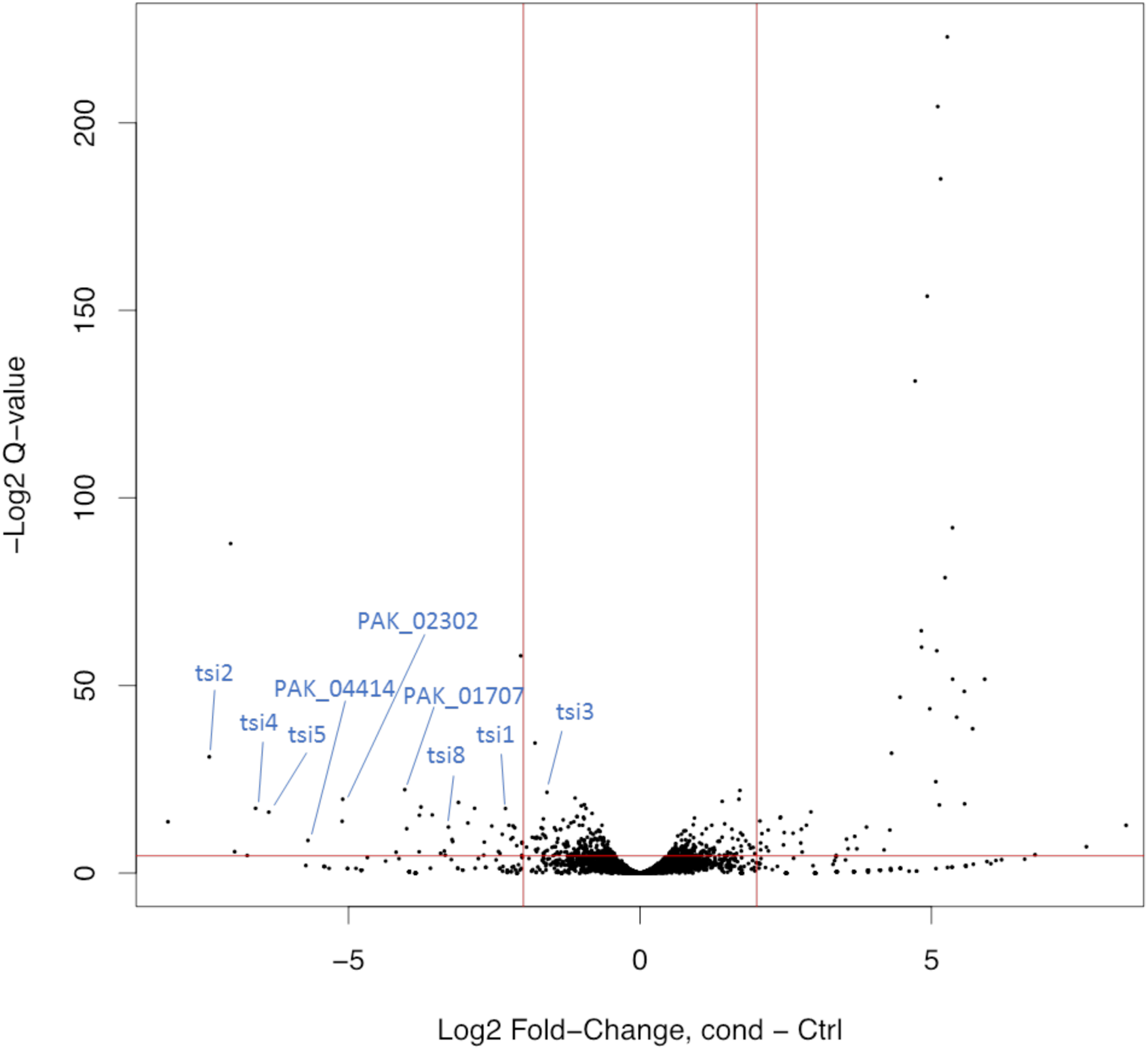
Volcano plot showing the spread of changes in abundance of TraDIS mutants for each *P. aeruginosa* gene during T6SS active compared to inactive conditions. Each black dot represents the comparative fold change of insertions for each gene. Red lines show the cut off criteria of 5% false discovery rate (horizontal) and a log_2_ fold change (Log_2_FC) of 2 (vertical). Immunity genes and putative immunity genes (as shown in Table 1) are shown in blue.

