## Supplementary information for "Discovery of a *Pseudomonas aeruginosa* Type VI secretion system toxin targeting bacterial protein synthesis using a global genomics approach"

### Legends for Supplementary Tables

**Supplementary Table 1.** Contains the full TraDIS data set (Tab 1), data above our threshold cut-off (-logFC >2 q<0.05) (Tab 2), and data from Tab 2 limited to genes <600 bp (Tab 3). See 'Generation of TraDIS sequencing libraries, sequencing and downstream analysis' below for column information.

**Supplementary Table 2.** Output of bioinformatic analyses of relevant bacteria possessing GatCAB, GlnS and AsnS.

### Supplementary Methods

#### TraDIS library generation

The *E. coli* donor strain was grown in LB supplemented with gentamicin (15 μg/mL) overnight at 37 °C and the recipient PAK strain was grown overnight at 37 °C in LB. Equivalent amounts of both strains were spread uniformly on separate LB agar plates and incubated overnight at 37 °C for *E. coli* and at 43 °C under humid conditions for the PAK recipient. The next day one *E. coli* donor plate was harvested and combined by extensive physical mixing on a fresh LB agar plate with one plate of harvested recipient PAK strain. Conjugation between the two strains was achieved by incubation of the high-density mixture of both strains at 37 °C for 2 hrs under humid conditions. The conjugation mix was then harvested, pelleted by centrifugation (10,000 *g*, 10 mins, 4 °C), and resuspended in LB. The resuspended cells were recovered onto large square (225 mm) Vogel-Bonner Media (VBM) (MgSO4.7H2O (8 mM), citric acid (anhydrous) (9.6 mM), $K_2HPO_4$ (1.7 mM), $NaNH_5PO_4.4H20$ (22.7 mM), pH 7) agar plates supplemented with gentamicin (60 μg/mL) and incubated for 16 hrs at 37 °C. The numbers of mutants obtained were estimated by counting a representative number of colonies across multiple plates. Mutants for each library background on plates were recovered as two separate pools (T6SS active and T6SS inactive), resuspended in LB, then pelleted by centrifugation (10,000 *g*, 10 mins, 4 °C), and then finally resuspended in LB plus glycerol (15% (v/v)) and stored at -80 °C. The protocol was repeated on a large scale until ~2 million mutants were obtained in each library background. For the TraDIS assay glycerol stocks of harvested PAKΔ*retS* or PAKΔ*retS*ΔH1 TraDIS libraries were combined at normalized cell density for each separate replicate (*i.e* two final pools in total) and spread onto large square (225 mm) VBM agar plates supplemented with gentamicin (60 µg/mL) and incubated for 16 hrs at 37 °C to facilitate T6SS delivery of toxins and subsequent killing/self-intoxication of mutants lacking immunity genes for the cognate toxin. Cells were then harvested into 5 mL LB and pelleted by centrifugation (10,000 *g*, 15 mins, 4 °C). Cell pellets were resuspended in 1.4 mL LB and 1 mL was retained for subsequent genomic DNA extraction (see 'TraDIS library genomic DNA extractions' section in the Methods of the main manuscript).

**Generation of TraDIS sequencing libraries, sequencing and downstream analysis**

PCR primers were designed for library construction and used for both the PAK libraries (5´: AATGATACGGCGACCACCGAGATCTACACACAGGAAACAGGACTCTAGAGG ATCACC and 3´: AATGATACGGCGACCACCGAGATCTACACCTTCTGTATGGAACG GGATGCG) and the sequencing TraDIS primers (5´: CAGCTTTCTTGTACACTAGA GACCGGGGACTTATCAG, and 3´: AAGCCTGCTTTCTAGAGACCGGGGACTTAT CAG). During library production, a post-ligation double digest with restriction enzymes AgeI and SgrAI was performed according to the manufacturer's instructions (New England Biolabs) to prevent amplification of plasmid background. The T6SS TraDIS sequencing was performed on a HiSeq2500 Illumina platform on the RAPID 50bp SE read setting. Reads were mapped onto the PAK genome (accession number: ERS195106), and comparisons were performed using the TraDIS Toolkit informatics package[4]. 10% of the 3´ end of each gene was discounted, and a 10 read minimum cut-off was used to be included in analysis. On average there was a unique transposon insertion site every 53 bp over the whole genome for each of the T6SS active and T6SS inactive backgrounds and, thus the genome was highly saturated in each library. The distribution of transposon insertions across the genome based upon the normalized transposon insertions in a H1-T6SS inactive library background, compared to the H1-T6SS active library background is shown in Extended Fig. 9. The resulting sequences of the T6SS TraDIS assays are available from the European Nucleotide archive (ENA) under study accession number PRJEB1597.

To pinpoint genes involved in protection of T6SS-mediated killing, EdgeR[5] was used to identify significant differences in read counts of genes in strains with (PAKΔ*retS*) and without (PAKΔ*retS*Δ*H1*) an active H1-T6SS. Then the trimmed mean of M values (TMM) normalization was used to account for differences between then libraries, and tagwise dispersion was estimated. Only genes exhibiting greater than 5 reads in both replicates of the conditions or control sets were examined for differences in the prevalence of mutants. Genes with zero read counts in the other condition were offset using the prior count function in EdgeR[5] so that fold changes could be estimated. P values were corrected for multiple testing using the Benjamini- Hochberg method, and genes with a corrected P value (Q value) of <0.05 (5% false discovery rate) and an absolute log2 fold change (log2FC) of >2 were considered significant (see Tab 2 in Supplementary Table 1). A list of 49 genes resulted having statistically significant decreased insertions in the T6SS active library PAKΔ*retS* compared to normalized values in the PAKΔ*retS*Δ*HI* library. These genes were then interrogated as potential immunities, based firstly on gene size (the known H1-T6SS associated immunity genes (*tsi1-6*) at the time of analysis are all less than 600bp, thus this was used as a guide to shorten the list to 29 genes) (see Tab 3 in Supplementary Table 1) and also on whether a protein upstream these genes appeared to have a predicted enzymatic or putative toxin function.

**Expression and purification of Tse8 used for activity measurements**

The pET41a::GST-TEV-Tse8 vector coding for *P. aeruginosa* Tse8 was obtained by FastCloning[6] using pET41a:GST-Tse8 (see Extended Table 1) as template. This construct was subcloned using the forward primer 5´-AACCTGTATTTTCAGGGCGGATCC ATCGAGGTCACCGAGGTTTCCATCG-3´ and reverse primer 5´-CCTGAAAATACAGG TTTTCGGTACCCAGATCTGGGCTGTCCATGTGCTGG-3´ in order to exchange the *Human Rhinovirus* (HRV) 3C cleavage site (LEVLFQ/GP) with a TEV protease cleavage site (ENLYFQ/G). The resulting construct includes (*i*) a 651-nucleotide sequence encoding a N-terminal GST tag, (*ii*) an 18-nucleotide sequence encoding a 6x histidine tag, (*iii*) a 45-nucleotide sequence encoding a S15 tag and a 21-nucleotide sequence encoding the optimal tobacco etch virus (TEV) protease cleavage site Glu-Asn-Leu-Tyr-Phe-Gln-Gly (Extended Fig. 5). For protein expression, *E. coli* BL21(DE3) cells were transformed with the pET41a::GST-TEV-Tse8 plasmid and grown in 2xYT (Yeast Extract Tryptone) medium (supplemented with 50 μg/ml kanamycin) at 37 ºC. When the culture reached an $OD_{600}$ value of 0.7, Tse8 expression was induced by adding 1 mM isopropyl β-D-1-thiogalactopyranoside (IPTG) and the temperature was dropped to 18°C. After 18 hr, cells were harvested and frozen for later use.

For protein purification, each 1 L pellet was resuspended in 50 ml of 50 mM Tris-HCl pH 8, 500 mM NaCl, 20 mM imidazole, 0.5 mM EDTA and 2 μL of benzonase endonuclease (without addition of protease inhibitors). Cells were then disrupted by sonication and the suspension was centrifuged for 40 mins at 56,000 *g*. The supernatant was filtered with a 0.2 μm syringe filter and subjected to immobilized metal affinity chromatography using a 1 ml HisTrap HP column (GE Healthcare), on a fast protein liquid chromatography system (ÄKTA FPLC; GE Healthcare) equilibrated with 5 ml of 50 mM Tris-HCl pH 8, 500 mM NaCl and 20 mM imidazole (buffer A). The column was washed with buffer A at 1 ml/min until no absorbance at 280 nm was detected. Elution was performed with a linear gradient between 0-

50% of 50 mM Tris-HCl pH 8, 500 mM NaCl and 500 mM imidazole in 30 mL and at 1 ml/min. Fractions containing GST-TEV-Tse8 fusion protein were pooled and protein concentration was measured. The cleavage of the GST-His-S15 tag was performed with TEV protease (1 mg per 10 milligrams of protein) overnight at 18 ºC in buffer 50 mM Tris–HCl pH 7.5, 2 mM DTT, at a protein concentration between 0.3-0.5 mg/mL. The cleaved Tse8, non-cleaved Tse8 and TEV protease were collected, filtered and applied onto a HisTrap HP column (5 ml; GE Healthcare) equilibrated with 25 ml of 50 mM Tris-HCl pH 7.5. The cleaved Tse8 was eluted in the flow-through and applied onto a Mono Q column of 5 mL (GE Healthcare) equilibrated with 25 ml of 50 mM Tris-HCl pH 7.5. The protein was eluted in a single step using 500 mM NaCl in 50 mM Tris-HCl pH 7.5. The Tse8 protein was dialyzed with 20 mM sodium phosphate buffer pH 7.6 and concentrated using Centricon centrifugal filter units of 30 kDa molecular mass cut-off (Millipore) to a final concentration of 5 mg/mL for enzymatic assays. The purity of the protein was verified by SDS-PAGE (Extended Fig. 5) and protein integrity was evaluated following desalting with stage-tip $C_4$ microcolumns (Zip-tip, Millipore) by electrospray ionization mass spectrometry (ESI-MS). The sampling cone energy was set at 35 V. The *m/z* data were then deconvoluted into MS-data using the MaxEnt software (MaxEnt Solutions Ltd, Cambridge, UK) with a resolution of the output mass of 0.5 Da/channel and Uniform Gaussian Damage Model at the half height of 0.5 Da. The analysis indicates that 90% of the protein sample corresponds to the expected Tse8 molecular weight (60,564 Da; Extended Data Fig. 5).

**Mass spectrometry analysis of Tse8/Pam enzymatic assays using epinecidin-1 as a substrate**

Samples were desalted and peptides were isolated using stage-tip $C_{18}$ microcolumns (Zip-tip, Millipore) and further resuspended in 0.1% formic acid prior to MS analysis. Peptide separation was performed on a nanoACQUITY UPLC System (Waters) on-line connected to an LTQ Orbitrap XL mass spectrometer (Thermo Electron). An aliquot of each sample was loaded onto a Symmetry 300 C18 UPLC Trap column (180 µm x 20 mm, 5 µm (Waters)). The precolumn was connected to a BEH130 $C_{18}$ column (75 µm x 200 mm, 1.7 µm (Waters), and equilibrated in 3% acetonitrile and 0.1% FA. Peptides were eluted directly into an LTQ Orbitrap XL mass spectrometer (Thermo Finnigan) through a nanoelectrospray capillary source (Proxeon Biosystems), at 300 nl/min and using a 120 mins linear gradient of 3-50% acetonitrile. The mass spectrometer automatically switched between MS and MS/MS acquisition in DDA mode. Full MS scan survey spectra (*m/z* 400-2000) were acquired in the orbitrap with mass resolution of 30,000 at *m/z* 400. After each survey scan, the six most intense ions above 1,000 counts were sequentially subjected to collision-induced dissociation (CID) in the linear ion trap. Precursors with charge states of 2 and 3 were specifically selected for CID. Peptides were excluded from further analysis during 60 s using the dynamic exclusion feature. RAW files were searched with the Mascot search engine (www.matrixscience.com) through Proteome Discoverer v1.4 (Thermo) against a FASTA database containing the protein and peptide sequences of interest, together with a *Pichia pastoris* database from Uniprot/Swissprot as a background. Search parameters were: 10 ppm peptide mass tolerance, 0.5 Da fragment mass tolerance, carbamydomethylation of cysteines as fixed modification, and oxidation of methionine, amidation and deamidation of protein C-terminus as variable modifications. Only highly reliable hits ($p<0.01$) were considered.

**Mass spectrometry analysis of Tse8 enzymatic assay using glutamine as substrate**

Overnight incubations were quenched by addition of 150 µL 20% acetonitrile (MeCN). Controls for the experiment were prepared by first adding MeCN to the reaction blank and subsequently adding enzyme. In order to determine LC-MS performance, 100 µM stock solutions of glutamine substrate in 2:3 water/MeCN were injected before the experimental samples. Quenched incubations and controls were shaken in the tubes for 30 mins at 4°C and 1,000 *g*. Next, samples were centrifuged for 30 mins at 4°C and 25,000 *g*. The resulting solutions were immediately injected in the LC-MS. Samples were measured with a UPLC system (Acquity, Waters Inc., Manchester, UK) coupled to a Time of Flight mass spectrometer (ToF MS, SYNAPT G2, Waters Inc.). A 2.1 x 100 mm, 1.7 µm BEH amide column (Waters Inc.), thermostated at 40 °C, was used to separate the analytes before entering the MS. Mobile phase solvent A (aqueous phase) consisted of 99.5% water, 0.5% formic acid and 20 mM ammonium formate while solvent B (organic phase) consisted of 29.5% water, 70% MeCN, 0.5% formic acid and 1 mM ammonium formate. In order to obtain a good separation of the analytes the following gradient was used: from 5% A to 50% A in 2.4 mins in curved gradient (#8, as defined by Waters), from 50% A to 99.9% A in 0.2 mins constant at 99.9% A for 1.2 mins, back to 5% A in 0.2 mins. The flow rate was 0.250 mL/min and the injection volume was 2 µL. The MS was operated in positive (ESI+) and negative (ESI-) electrospray ionization in full scan mode. The cone voltage was 25 V and capillary voltage was 250 V for ESI+ and 500 V for ESI-. Source temperature was set to 120 °C and capillary temperature to 450 °C. The flow of the cone and desolvation gas (both nitrogen) were set to 5 L/h and 600 L/h, respectively. A 2 ng/mL leucine-enkephalin solution in water/acetonitrile/formic acid (49.9/50/0.1% (v/v/v)) was infused at 10 µL/min and used for a lock mass which was measured each 36 seconds for 0.5 seconds. Spectral peaks were automatically corrected for deviations in the lock mass.

**Bioinformatics analysis of prokaryotic organisms encoding AsnS and GlnS**

*Escherichia coli* AsnS and GlnS protein sequences were interrogated against the National Center for Biotechnology Information (NCBI) collection of non-redundant protein sequences of bacteria and archaea (*non-redundant Microbial proteins*, update: 2017/11/29) using the *pBLAST* search engine. The search was further restricted for *non-redundant RefSeq* *proteins*, with a 20,000-hit limit, the BLOSUM62 matrix scoring function and an *Expect threshold value* (*E-value*) of $1e^{-5}$. Hits were selected if sequence identity was above 50% with respect to the query sequences and those associated with bacterial species *Agrobacterium tumefaciens, Escherichia coli* and *Pseudomonas aeruginosa* were extracted (Supplementary Table 2).

**Bioinformatics analysis of prokaryotic organisms predicted to encode the amidotransferase GatCAB complex**

A none-exhaustive search for organisms encoding GatCAB was carried out using the National Center for Biotechnology Information (NCBI) database. *Pseudomonas aeruginosa* GatA and GatB protein sequences were interrogated against the NCBI collection of non-redundant protein sequences of bacteria and archaea (*non-redundant Microbial proteins*, update: 2017/11/29) using the *pBLAST* search engine. The search was further restricted for *non-redundant RefSeq proteins*, with a 20,000-hit limit, BLOSUM62 matrix scoring function and an *Expect threshold value* (*E-value*) of 1e-5. Hits were selected if annotated as Asp-tRNA(Asn)/Glu-tRNA(Gln) amidotransferase subunits, and the results for *Agrobacterium tumefaciens, Escherichia coli* and *Pseudomonas aeruginosa* were extracted (Supplementary Table 2).
